# From Substrate to Short Chain Fatty Acids: An *In Vitro* Investigation of Bacterial Dynamics in the Porcine Colon using the Modified Hohenheim Gas Test

**DOI:** 10.64898/2026.11.26.769249

**Authors:** Ina Klawes, Marvin Suhr, Daniel Berghaus, Corinna Bang, Stéphanie Céline Hornburg

**Author notes:** Corresponding author: Stéphanie Céline Hornburg (.

## Abstract

**Background:** Research on the development of pig colonic bacteria under *in vitro* conditions is limited. In this *in vitro* study, the modified Hohenheim Gas Test (modHGT) was used to improve the understanding of fermentation processes in the pig colon with respect to the underlying microbial community, in particular how different substrates modulate the microbial community structure during fermentation.

**Methods:** Gas formation, short chain fatty acid (SCFA) production by HPLC, bacterial load by qPCR and bacterial community structure by 16S rDNA amplicon sequencing were assessed at 2, 4, 8 and 24 hours during the fermentation of wheat straw, inulin, sugar beet pulp and potato starch using pig feces as microbial inoculum.

**Results:** The dominant phyla (Firmicutes, Bacteroidota, Proteobacteria) and genera observed were consistent with previous *in vivo* studies, but differences in abundance were found. Microbial activity varied significantly (*p* < 0.05) between substrates and time points, influencing gas formation and SCFA profiles (*p* < 0.05). Rapid shifts in bacterial communities within the first 2 hours highlight the strong substrate influence on fermentation dynamics.

**Conclusions:** The modHGT is a valuable tool for advancing our understanding of the porcine gut microbiota. This method is accessible, requires minimal effort and is in line with animal welfare considerations. It has great potential to provide insights into functional aspects of the gut microbiota and host interactions in different (feeding) situations. Metabolic interdependencies, such as cross-feeding, influence microbial community structure and SCFA profiles. One approach to standardization could be the use of standardized microbiome reference communities, similar to human gut microbiota research. Future research needs to investigate the consequences of altering bacterial composition and its physiological functions and consequences.

**Highlights:**

- *In vitro* pig gut fermentation offers an alternative to feeding trials
- *In vitro* gut microbiota resembles in vivo but differs in relative abundance
- Gut microbiota, gas and SCFA dynamics vary with fermented substrate
- Bacterial composition shifts rapidly depending on the fermented substrate
- Standardized microbiome references may enhance pig gut microbiota research

## 1. Introduction

Understanding microbial community dynamics in the gastrointestinal tract is crucial for decision making in pig nutrition. The gastrointestinal tract of mammals hosts a diverse and dynamic microbial community that is crucial for digestive processes (FLINT ET AL. 2012). A key component of porcine intestinal health is fermentation, during which short-chain fatty acids (SCFA) such as acetic, propionic, and butyric acid, are produced (BUGAUT 1987). These fermentation products serve not only as energy sources for the host but also play essential roles in maintaining gut barrier function and modulating immune responses (WANG ET AL. 2022; SHIN ET AL. 2019; PATIL ET AL. 2020; LEE ET AL. 2023). Moreover, the ratio of these volatile fatty acids, which serves as a comparative measure of fermentation processes, is strongly influenced by the substrates available in the large intestine. Therefore, dietary composition is one of the most important factors affecting colonic fermentation processes (FRIEND ET AL. 1963; KENNELLY ET AL. 1981; BACH KNUDSEN AND HANSEN 1991; PATIL ET AL. 2020). Feeding trials have long been considered the gold standard for evaluating the effects of raw materials and diets on animals (KIRCHGEßNER 1980). However, these trials are time-consuming and often require long observation periods to determine effects. As a result, the delay in obtaining results can make timely decisions on feeding strategies difficult. In addition, in Europe animal experiments are regulated under European law and must comply with the Animal Welfare Act. In 2020, approximately 56,094 farm animals were used for scientific purposes in Germany, representing about 2.2% of all laboratory animals (TREUE ET AL. 2022). The Farm Animal Welfare Committee (FAWC, 1993) defines animal welfare through the “Five Freedoms”: freedom from hunger and thirst, discomfort, pain, fear, and the ability to exhibit natural behaviors. In feeding trials, stress from dietary changes or handling can affect both, welfare and experimental outcomes. Additionally, genetic and environmental factors can limit reproducibility and applicability across different settings. In response to these challenges, the German government has increasingly promoted the use of non-invasive methods, such as *in vitro* techniques and computer-based models, to reduce *in vivo* trials (BMEL 2024). Among these, *in vitro* approaches like the modified Hohenheim gas test (modHGT) offer rapid and reliable tools for assessing dietary effects on intestinal fermentation parameters (RINK ET AL. 2011). Yet, little attention has been paid to the *in vitro* community dynamics using pig feces as microbial inoculum. The main objective of this study was to enhance the understanding of fermentation processes in relation to the underlying microbial community, with a particular focus on how different substrates influence community structure during *in vitro* fermentation using a modified Hohenheim gas test. Gas formation was monitored at 2, 4, 8 and 24 hours of fermentation. Accordingly, SCFA-profiles, bacterial load, and bacterial community dynamics were assessed via 16S rDNA amplicon sequencing. Four substrates, inulin, potato starch, sugar beet pulp, and wheat straw were selected for their diverse fermentation characteristics. These substrates represent different carbohydrate compositions, allowing insights into how substrates influence microbial activity and community dynamics.

## 2. Material and methods

In this study, inoculum from porcine fecal samples were incubated with four different substrates (inulin, potato starch, sugar beet pulp and wheat straw) in a modified Hohenheim Gas Test. Gas formation was monitored for 24 hours and gas formation was measured after 2, 4, 8 and 24 hours. Simultaneously, samples were taken for SCFA analysis via High-Performance Liquid Chromatography (HPLC), and bacterial 16S rDNA amplicon sequencing and qPCR analysis. The experimental scheme is outlined in Figure 1.

**Figure 1:**
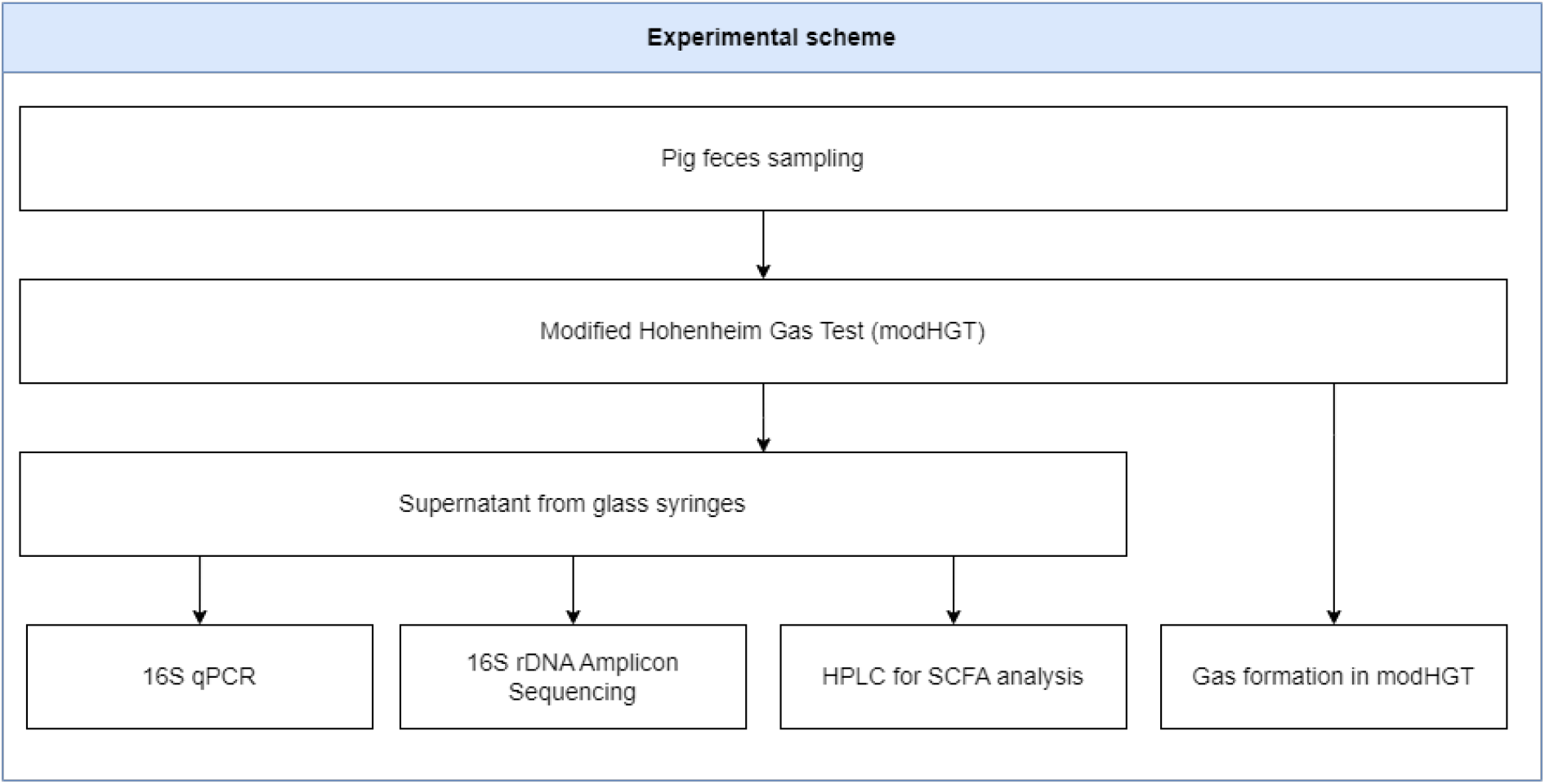
Schematic representation of the modified Hohenheim Gas Test (modHGT) with prior pig feces sampling. Supernatants of glass syringes were analyzed to determine the bacterial community composition and bacterial load, as well as SCFA production. Gas formation was visually monitored during the incubation experiment. Created with draw.io 2024

### 2.1. Animals and sampling

The animals used as donor pigs for this study were kept farm animals and housed in accordance with EU and national law (Federal Ministry of Justice. 2006b) and the ‘German Animal Welfare Ordinance on Farm Animal Husbandry’ (Federal Ministry of Justice 2006a) in group pens on a conventional farm (Gut Hülsenberg GmbH, Wahlstedt, Germany). To minimize animal-specific variation in fecal bacterial composition and potential dietary influences, feces samples were collected from six female 18-week-old growing pigs (Danbred × Duroc) within a body weight range from 72 to 94 kg. All animals were healthy, raised under the same housing conditions and were fed a standard diet formulated to meet the requirement for an average daily gain of 950 g (GfE 2008) based on barley, wheat, maize, and soybean meal (Table 1), provided three times a day (morning 7:00, midday 11:00, evening 16:00) in liquid form Drinking water was available continuously via nipple drinkers. Analyzed nutrient composition of the diet of the donor animals is presented in Table 2.

**Table 1:** Formulation of the diet in grower phase by raw material and dry matter content (%)

| Ingredient | DM | % |
|---|---|---|
| Barley | 880 | 36.00 |
| Wheat | 880 | 28.00 |
| Maize | 880 | 20.00 |
| Soybean oil | 990 | 0.50 |
| Soybean meal | 880 | 9.50 |
| Mineral and Vitamin premix<sup>1</sup> | 980 | 3.00 |
| Lignocellulose | 960 | 0.50 |
| Acid blend<sup>2</sup> | 1000 | 0.50 |
| Oat bran | 910 | 2.00 |

1: Vitamin A, D, E, iron, zinc, manganese, copper, iodine, selenium, 6-phytase, calcium carbonate, sodium chloride, monocalcium phosphate, magnesium oxide (NATUPIG M 120) provided by H. Wilhelm Schaumann GmbH, Pinneberg, Germany
2: Formic acid, lactic acid, propionic acid, provided by H. Wilhelm Schaumann GmbH, Pinneberg, Germany (SCHAUMACID F)

**Table 2:** Analyzed nutrient composition of the diet in grower phase (as fed)

| Nutrient | % |
|---|---|
| Metabolizable Energy | 12.96 |
| Dry Matter | 88.52 |
| Crude Protein | 14.37 |
| Crude Fiber | 4.35 |
| Neutral Detergent Fiber | 14.99 |
| Lysin | 0.91 |
| Calcium | 0.60 |
| Phosphorus | 0.35 |

At least two weeks prior to sampling, the animals were not treated with any medication, in particular antibiotics, in order to avoid any impact on the composition and activity of the intestinal microbial community. Sampling took place during the grower phase, three weeks after the transition from starter to grower feed, in a separated barn compartment in the absence of non-donor animals before morning feeding. Feces were collected per rectum during spontaneous defecation using plastic bags. To maintain anaerobic conditions after sampling, samples were immediately placed in plastic containers, sealed and transported to the on-site laboratory in thermoboxes (Thermohauser GmbH, Uhingen, Germany) and immediately prepared for incubation to minimize any changes in the bacterial community.

### 2.2. Incubated substrates

Four different fermentable fibers (wheat straw, inulin, potato starch, and sugar beet pulp) were used as incubation substrates (Table 3). The four substrates were selected to represent potentially fermentable fibers from different carbohydrate classes and potential components of pig diets. Wheat straw is a by-product of the production and harvesting of wheat (*Triticum spp.).* It consists primarily of the dry stalks of the wheat plant which are composed of cellulose, hemicellulose, and lignin (ZHANG ET AL. 2022). Inulin is a fructan polysaccharide found in various plant roots and tubers such as chicory and artichokes. It consists of chains of fructose molecules linked together by β-(2,1)-glycosidic bonds (QIN ET AL. 2023). Potato starch is a polysaccharide consisting of glucose linked by α-1,4-glycosidic bonds and is the main component of potato tuber (BERTOFT AND BLENNOW 2009). Sugar beet pulp is a mixture of polysaccharides including cellulose, hemicellulose, and pectin which is a by-product of sugar beet processing in the sugar and ethanol industries (WALDRON 2007-2009). All substrates were obtained from commercial sources and were air-dried and ground to pass a 1mm sieve (ISO 6498). Determination of dry matter (DM), crude ash (XA), crude protein (XP), crude fiber (XF), neutral detergent fiber (aNDFom), and acid detergent fiber (ADFom) was performed according to VDLUFA procedures (Verband Deutscher Landwirtschaftlicher Untersuchungsund Forschungsanstalten (VDLUFA) 2007)

**Table 3:**
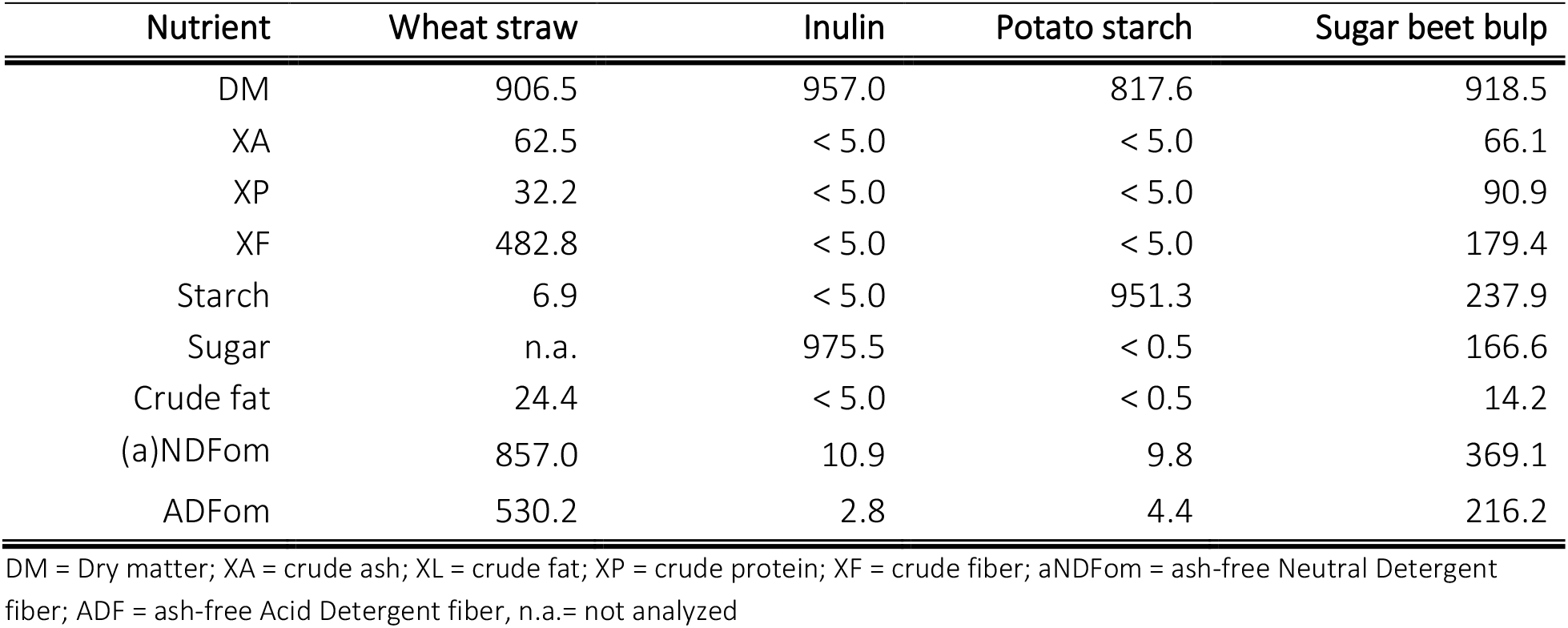
Chemical composition of substrates used for incubation (as fed)

| Nutrient | Wheat straw | Inulin | Potato starch | Sugar beet bulp |
|---|---|---|---|---|
| DM | 906.5 | 957.0 | 817.6 | 918.5 |
| XA | 62.5 | < 5.0 | < 5.0 | 66.1 |
| XP | 32.2 | < 5.0 | < 5.0 | 90.9 |
| XF | 482.8 | < 5.0 | < 5.0 | 179.4 |
| Starch | 6.9 | < 5.0 | 951.3 | 237.9 |
| Sugar | n.a. | 975.5 | < 0.5 | 166.6 |
| Crude fat | 24.4 | < 5.0 | < 0.5 | 14.2 |
| (a)NDFom | 857.0 | 10.9 | 9.8 | 369.1 |
| ADFom | 530.2 | 2.8 | 4.4 | 216.2 |

DM = Dry matter; XA = crude ash; XL = crude fat; XP = crude protein; XF = crude fiber; aNDFom = ash-free Neutral Detergent fiber; ADF = ash-free Acid Detergent fiber, n.a.= not analyzed

### 2.3. *In-vitro* fermentation

To evaluate gas formation as a metabolic end product of microbial fermentation processes of the four substrates, a modHGT was conducted. Therefore, pig feces were used as microbial inoculum in a 1:60 dilution (on DM basis) with buffer solution as described by RINK ET AL. (2011) with slight adjustments. Briefly, the buffer solution was prepared according to BUSKE ET AL. (2000) before sampling and kept at a constant 39°C while stirring with a magnetic stirrer to maintain homogeneity. Then, equal amounts of feces (60 g, wet weight) from each animal were pooled with the same amount of the prewarmed buffer solution using a conventional kitchen blender (Thermomix®, Vorwerk SE & Co. KG, Wuppertal, Germany), which was fitted with a modified rubber plug in the lid, through which CO2 was administered via a tube to maintain anaerobic conditions. After blending for 150 sec, 240 g of this premix was transferred to a CO2-flushed plastic bottle and diluted with 1560 g of the buffer solution to obtain the 1:60 feces-buffer dilution. Thereafter, the feces-buffer dilution was filtered through a clean cheese cloth into a Woulff bottle under a constant stream of CO2. Then, trace element, reduction and indicator solution were added to the feces-buffer dilution, the Woulff bottle was fitted with an automatic dispenser and placed in a water bath at 39°C with a magnetic stirrer to maintain a homogeneous inoculum. Glass syringes with a volume of 150 ml (100 ml calibrated, measuring dial in 0.5 ml intervals) were filled with 200 mg of substrate the day before fermentation, preheated overnight at 39°C, filled with 30 ml (± 1 ml) inoculum each and the initial liquid volume (V0) was documented. The syringes were then placed in an incubation chamber at 39°C with continuous mixing at one rotation per minute for 24 hours. After 2, 4, 8, and 24 hours, gas formation was determined by stopping the rotor and reading the syringes individually. If the gas volume risked exceeding the calibrated 100 ml scale during incubation, gas production was recorded at 8 hours, gas was released, and the inoculum volume was noted. Syringes with inoculum but without added substrate were used as blanks to correct for the gas volume attributable to the inoculum itself. For subsequent SCFA and microbiome analysis, two syringes per treatment were removed at 2, 4, and 8 hours. At 24 hours, two of the remaining four syringes per treatment were used for SCFA and microbiome analysis and the other two only for gas readings, resulting in a total of 50 syringes. After removal, the contents of the syringes were immediately filtered through a fine mesh and aliquots of the supernatant were taken and stored at –20 °C until further processing for SCFA analysis or at –80°C until DNA extraction for microbiome analysis.

### 2.4. Gas formation

Net gas formation was measured in calibrated syringes according to MENKE ET AL. (1997) and corrected to 200 mg DM of the test substrate with following equations:

Gas formation (GF, ml) without degasification:

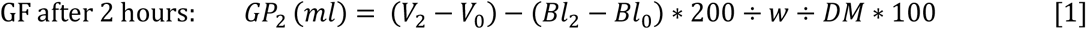

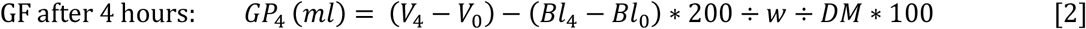

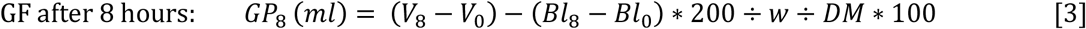

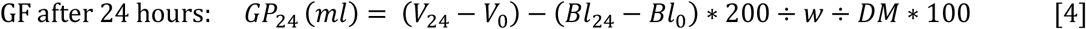

GF with degasification after 8 h of incubation:

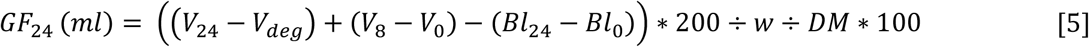

With:

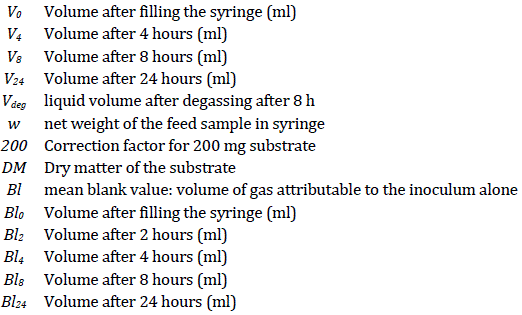

According to MENKE ET AL. (1979) the blank GF was subtracted from every measurement to calculate net GF.

### 2.5. Short chain fatty acid analysis

SCFA, including acetic acid, propionic acid, butyric acid, lactic acid and the total SCFA (sum of acetic acid, propionic acid, isobutyric acid, butyric acid, isovaleric acid, valeric acid, and lactic acid), were quantified by HPLC at ISF Schaumann Forschung GmbH (Wahlstedt, Germany) using appropriate internal standards. Sample preparation and the analysis of SCFA were performed as described by WISCHER ET AL. (2013). Briefly, samples were thawed, stirred, and centrifuged at 4,000 g for 5 min. One ml of the supernatant was transferred into an Erlenmeyer flask and mixed with 0.1 ml of an internal standard solution (80 mM 2-methylvaleric acid in 50% formic acid). The samples were then frozen in an alcohol bath at −20°C and vacuum distilled. The distillates were analyzed in duplicate. For extraction, 5 g of each sample was mixed with 5 g of distilled water and stirred intensively for 10 minutes. After purification by centrifugation and filtration, the sulfuric acid concentration was adjusted to match the mobile phase (0.02 n sulfuric acid). Chromatographic separation was performed on an Aminex HPX-87H column (300 × 7.8 mm, Bio-Rad, Cat. No.: 125-0140) with a corresponding pre-column (Micro-Guard Cartridge HPX-87H, Bio-Rad, Cat. No.: 125-0129). The sample volume injected was 50 µl. Separation was achieved using 0.02 n sulfuric acid as the mobile phase under isocratic conditions with a flow rate of 0.6 ml/min at a temperature of 30–35°C. Detection was performed using a UV detector (Smartline 2500) at 210 nm and an RI detector (Smartline 2300). The system was equipped with a column thermostat (Jetstream 2, Bio-Rad Laboratories, California, USA). For quantification, external standards of pure organic acids were used. Pre-treated extracts were diluted to ensure the expected values fell within the calibrated range. The content of organic acids was calculated by considering the extraction fluid-to-sample ratio and the dilution factor of the extract. Individual SCFA values were corrected for those present in blanks. Total SCFA was calculated as the sum of acetic acid, propionic acid, isobutyric acid, butyric acid, isovaleric acid, valeric acid, and lactic acid.

### 2.6. Bacterial load quantification via qPCR

Prior to qPCR analysis, concentration and purity of DNA samples were determined by using a Spectrophotometer (Thermo Scientific NanoDrop One, Thermo Fisher Scientific, Waltham, USA). A reference solution was used to establish the baseline spectrum before samples were applied to the spectrophotometer cuvette. Following sample thawing and homogenization, 2 µl of each sample was pipetted into the cuvette, and the absorbance at 260 nm (A260) and 280 nm (A280) was measured. The A260/A280 ratio was utilized to assess DNA purity, aiming for an ideal ratio of approximately 1.8 – 2.2 (Thermo Scientific, 2016). DNA concentration was calculated in ng/µl. Quantitative real-time PCR to determine total bacterial quantification over time was performed using a Thermocycler Rotor-Gene Q (QIAGEN N.V., Hilden, Germany) associated with the Rotor-Gene Q Series Software (version 2.3.5). ZymoTaq^TM^ qPCR PreMix was used according to the manufacturer’s instructions (Zymo Research, Freiburg, Germany). The total number of 16S rRNA genome copy numbers (GCN) per ml of incubation liquid supernatant was determined qPCR with specific primers for V3 – V4 region (341 Forward: CCTACGGGNGGCWGCAG; 785 Revers: GACTACHVGGGTATCTAATCC10:54) and 2 ng DNA in a total volume of 10 μl and expressed as 16S rRNA gene copies per ml of culture. The qPCR cycling protocol begins with an initial denaturation at 95°C for 10 minutes, followed by 40 cycles of denaturation at 95°C for 30 seconds, annealing at 55°C for 30 seconds, and extension at 72°C for 60 seconds. A final extension step is carried out at 72°C for 7 minutes, with a holding step at 4°C for at least 4 minutes. Standard curves were generated using serial dilutions of extracted DNA of bacterial cell cultures of known quantity as based on cell counts. *E. Coli* strain JM109 was used to generate calibration standards in duplicate according to the manufacturer’s instruction (E2006; Femto Bacterial DNA Quantification Kit, Zymo Research, Freiburg, Germany)

### 2.7. PCR and 16S rRNA gene amplicon sequencing

A total of 250 mg of pre-homogenized fermentation supernatant was mixed with 1 ml of InhibitEx buffer in 0.70 mm Garnet Bead tubes. Bacterial cell lysis was performed using a SpeedMill PLUS (Analytik Jena GmbH, Jena, DE) at high speed for 45 seconds. The lysate was then incubated for 5 minutes at 550 rpm and 95 °C. Following incubation, lysed cells were pelleted by centrifugation at 20.000 rcf for 1 minute. The total bacterial DNA extracted from incubation supernatant was subsequently isolated using 200 μl of the supernatant as input for the QIAamp® DNA Fast Stool Mini Kit (Qiagen, USA, Cat. no. 51604), following the manufacturer’s instructions on a QIAcube® automation platform (Qiagen, USA). Additionally, 12 extraction blanks were included to monitor potential contamination during DNA preparation. For sequencing, a one-step polymerase chain reaction (PCR) amplification of the V3–V4 region of the 16S rRNA gene was performed using the forward primer 341F ‘CCTACGGGAGGCAGCAG’ and reverse primer 805R ‘GGACTACHVGGGTWTCTAAT’ (KLINDWORTH ET AL. 2013), employing a dual-barcoding approach. The final reaction volume of 26 μl included 5.0 μl of 5X Phusion HF buffer, 0.5 μl of dNTPs (10 mM), 0.3 μl of Phusion Hot Start II Polymerase (2 U/μl; Thermo Fisher Scientific), 9.2 μl of H2O, 4 μl of each primer (100 μM), and 3 μl of microbial DNA template. Each PCR reaction plate contained a negative control with nuclease-free water to account for contamination, as well as a reaction with a mock community (ZymoBIOMICS Microbial Community Standard, Cat. no. D6305) as a positive control, consisting of eight bacterial isolates with defined abundances to verify adequate performance. The PCR conditions included an initial denaturation at 98 °C for 30 seconds, followed by 30 cycles of 98 °C for 9 seconds, 55 °C for 1 minute (annealing), and 72 °C for 1.5 minutes (extension). A final extension was performed at 72 °C for 10 minutes. The expected fragment size of 550 bp was confirmed via gel electrophoresis on a 2.0% agarose gel. Sequencing of the V3–V4 region of the 16S rRNA gene amplicons was performed using 54 samples. These included amplicons derived from the *in vitro* fermentation of different substrates with pig feces as a microbial inoculum, pig feces alone, as well as positive and negative controls. The sequencing was conducted at the Institute of Clinical Molecular Biology (IKMB) at Kiel University. An Illumina MiSeq platform (Illumina Inc., San Diego, CA, USA) was used, employing the MiSeq Reagent Kit v3. The procedure followed the manufacturer’s specifications and the protocol described by TRAUTMANN ET AL. (2020). This resulted in 2 × 300 bp paired-end reads. The raw sequences generated during the study are stored in the NCBI Sequence Read Archive (SRA) and can be accessed via the BioProjectID PRJNA1272849.

### 2.8. Bioinformatics

Demultiplexed reads, which were free from sequencing adapters, were provided by the sequencing facility as FASTQ files. The quality of these reads was assessed using FastQC (ANDREWS 2010) and MultiQC (EWELS ET AL. 2016). FASTQ files were then imported into Quantitative Insights Into Microbial Ecology 2 (Qiime2 2021.2.0; BOLYEN ET AL. (2019), where the following processing steps were performed. The Cutadapt plugin (MARTIN 2011) was used to trim any residual primers and/or spacers. To denoise, filter for low-quality reads and chimeras, and merge paired-end reads, DADA2 plugin (CALLAHAN ET AL. 2016) was employed with truncation parameters of 258 for forward reads, 222 for reverse reads. This process resulted in a feature table based on the Amplicon Sequence Variants (ASVs) and their corresponding representative sequences. Features occurring in less than two samples or with an overall frequency of less than 16 were discarded. To assign taxonomy to ASVs, a naïve Bayes classifier was trained against a SILVA 99% database (release 138). Regions of interest were extracted from full-length sequences using the V3–V4 primers and the feature-classifier plugin. Custom reference sequences were generated by extracting reads from the SILVA database. Features classified as Cyanobacteria, which were assumed to originate from chloroplasts, as well as those of mitochondrial origin, were excluded from further analysis. Only ASVs classified as bacteria and identified at least to the phylum level were retained. A phylogenetic tree was generated using a *de novo* approach by performing a multiple sequence alignment with a fast Fourier transform through the QIIME-phylogeny plugin. Mock community samples were separated from the dataset and analysed to ensure accurate expected abundance, as per the manufacturer’s instructions. Subsequent analyses were conducted using RStudio (version 4.3.1; 2023-06-16 ucrt) and the R package phyloseq (MCMURDIE AND HOLMES 2013). For alpha diversity analysis, richness was estimated based on observed features and Shannon diversity, with all parameters calculated at the genus level. To assess beta-diversity and compare the number of shared taxa among samples, Bray-Curtis distance was calculated using the R package vegan (ORKANSEN ET AL. 2008) and visualized using Non-metric Multidimensional Scaling (NMDS).

### 2.9. Model fitting and statistical analysis

GraphPad Prism4 software (2003) was used to fit sigmoidal curve for gas formation. Gompertz model equation fitting degree (R^2^) for gas data was > 0.98. The Gompertz Equation for fitting bacterial growth data (sigmoidal) was:

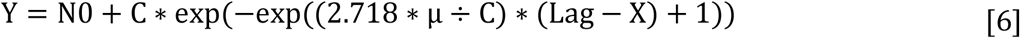

With:

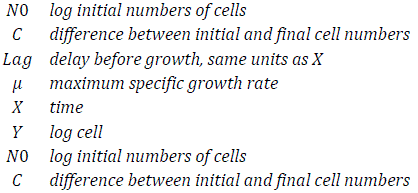

The statistical software R (version 4.3.1; 2023-06-16 ucrt) was used to evaluate gas production, SCFA formation, bacterial load (genome copy numbers) and bacterial community structure. The data evaluation started with the definition of an appropriate linear model. The model included the factors substrate and incubation time as well as their interaction term. The residuals were assumed to be normally distributed and to be homoscedastic. These assumptions are based on a graphical residual analysis. The linear model for gas formation, SCFA concentration and alpha diversity was:

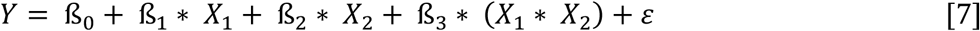

With:

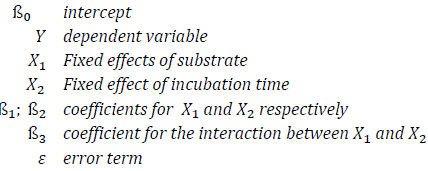

Based on this model, an analysis of variances (ANOVA) was conducted, followed by multiple contrast tests (BRETZ ET AL. 2011; HOTHORN ET AL. 2008) in order to compare the several levels of the influence factors, respectively.

To determine whether the bacterial community structure differed significantly between the samples, PERMANOVA was performed based on Bray–Curtis distances using the R “vegan” package (Oksanen et al., 2013) with 1000 permutations. A *p-value (corrected* with Bonferroni*) ≤0.05* was considered statistically significant for all analyses.

## 3. Results

### 3.1. Substrate fermentation determines gas formation, SCFA production and 16S rRNA gene copy numbers

#### 3.1.1. Gas formation during 24 h incubation of different substrates

Gas formation during the 24-hour fermentation of the four substrates (potato starch, inulin, sugar beet pulp, wheat straw) by fecal inoculum from growing pigs was modeled using the Gompertz function (Figure 2). Statistical analysis (see supplementary material file 1: gas formation results and statistics) revealed significant differences in gas formation among all substrates, except for inulin and sugar beet pulp, for which no significant difference was observed after 24 hours of incubation. The gas formation profiles varied significantly among the four substrates tested. To isolate the gas formation attributable solely to the inoculum, glass syringes containing only the inoculum (without additional substrates) were incubated. Minimal gas formation was observed under these conditions, with a total of 4 ml (*p* < 0.001) produced over the 24-hour incubation period. For potato starch, a lag phase was observed between 0 and 4 hours of incubation, during which gas formation was negligible. Following this period, a rapid increase in gas formation occurred, peaking at 8 hours. After reaching this peak, the rate of gas formation slowed down but continued until the 12-hour mark. Gas formation remained minimal beyond this point, leading to the highest final total gas formation of 63 ml (*p* < 0.001) by the end of the 24-hour incubation period. Inulin demonstrated a shorter lag phase (0 – 2 hours) compared to potato starch. Similar to potato starch, gas formation increased rapidly following this brief lag phase, reaching its peak at 5 hours of incubation. From 5 to 8 hours, gas formation decelerated, and thereafter, gas formation remained low, resulting in a final total of 60 ml after 24 hours. Microbial gas formation using sugar beet pulp as a substrate followed a continuous trend without a lag phase and without reaching a plateau within the observation period, despite similar total gas formation as for inulin after 24 hours (59.65 ml; *p* < 0.05). Straw showed the lowest gas formation among the substrates (*p* < 0.001). Rather than exhibiting a distinct lag phase, gas formation increased slowly and steadily over the entire incubation period, indicating continuous but low microbial activity, and resulting in the lowest cumulative gas formation after 24 hours.

**Figure 2:**
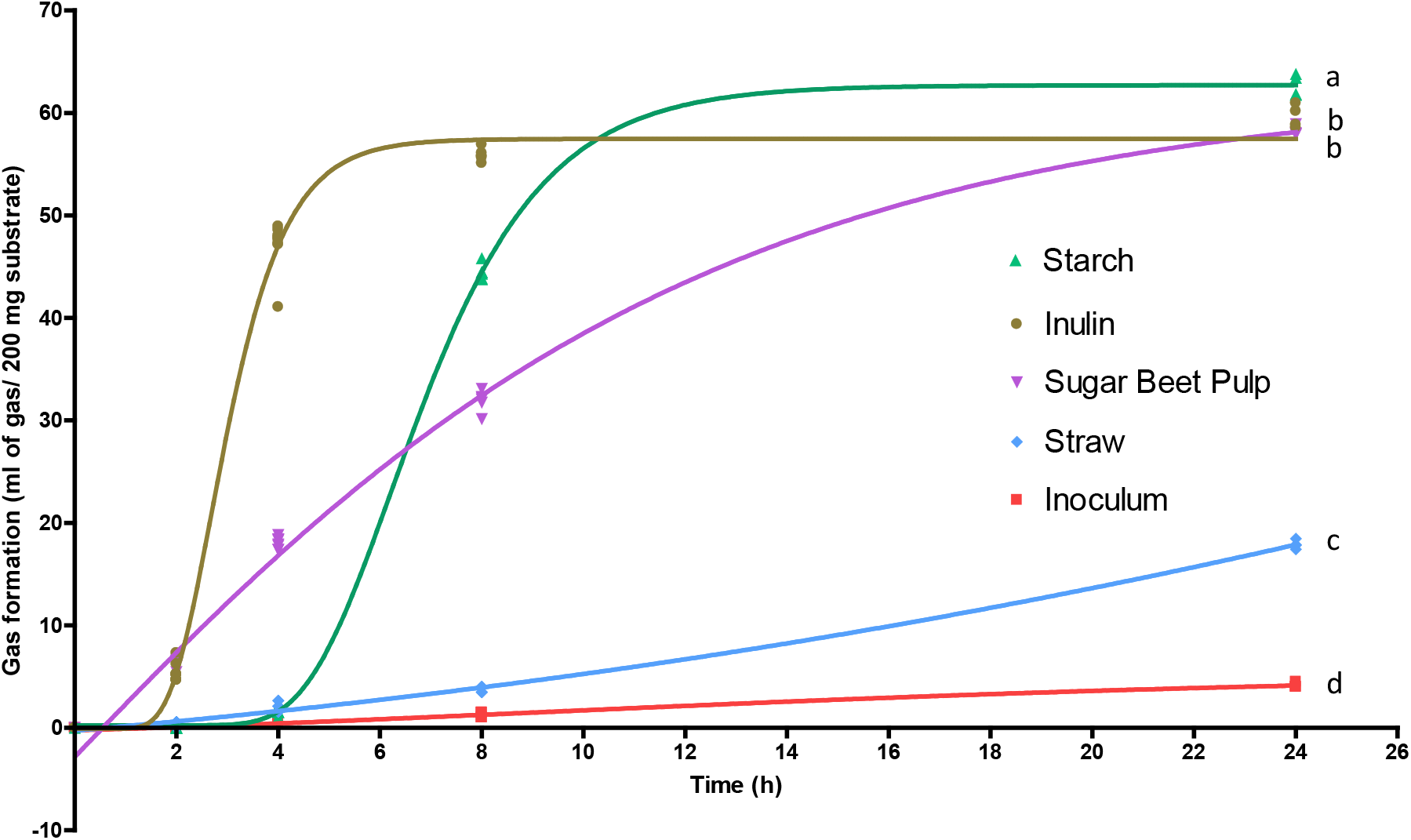
Gas formation during 24 hour-fermentation of four substrates (potato starch = green; inulin = ocher; sugar beet pulp = violet, wheat straw = blue) and fecal inoculum of growing pigs as a control (= red) was modelled using the Gompertz Function with the abscissa representing individual time points (2, 4, 8, 24 h), and the ordinate indicating gas formation in ml gas/ 200 mg substrate. R² for individual Gompertz models was ≥ 0.99; n = 4. Adjusted R² for ANOVA was 0.90. Differences were considered significant at p < 0.05. Substrates with different superscripts differ significantly.

#### 3.1.2. SCFA-Production during 24 hours incubation of different substrates

The analysis of SCFA production across different substrates over incubation periods of 2, 4, 8, and 24 hours revealed significant variations in the concentrations of the analyzed metabolites (see supplementary file 2: SCFA statistics). All concentrations are reported per 200 mg DM of substrate (Table 4). The total SCFA concentration for potato starch increased from 41.9 mg at 2 hours to 153.6 mg at 24 hours. Acetic acid concentrations notably rose from 17.6 mg to 70.7 mg over the same period. Butyric acid and propionic acid also increased, reaching 26.8 mg and 39.7 mg, respectively, at 24 hours. The production of SCFA from potato starch exhibited a progressive pattern, with acetic acid being the predominant SCFA throughout the incubation period. Inulin demonstrated the highest initial SCFA production, with a total concentration of 93.7 mg at 2 hours (*p* < 0.05), increasing to 166.8 mg at 24 hours.

**Table 4:** Production of short-chain fatty acids [mg per 200 mg dry matter of substrate] determined by HPLC after 2, 4, 8, and 24 hours of *in vitro* fermentation using the modified Hohenheim Gas Test (modHGT) with pig fecal inoculum and different substrates (wheat straw, inulin, sugar beet pulp, and potato starch).

| Substrate | t (h) | Total SCFA |  | Acetic Acid |  | Butyric Acid |  | Propionic Acid |  | Lactic acid |  |
|---|---|---|---|---|---|---|---|---|---|---|---|
| Inoculum | 2 | 35.3 | ± 0.4 | 15.0 | ± 0.2 | 6.3 | ± 0.1 | 9.3 | ± 0.2 | 0.2 | ± 0.1 |
|  | 4 | 40.3 | ± 0.5 | 17.8 | ± 0.8 | 7.1 | ± 0.1 | 10.6 | ± 0.1 | 0 | 0 |
|  | 8 | 46.6 | ± 0.1 | 21.7 | ± 1.0 | 7.7 | ± 0.0 | 11.6 | ± 0.0 | 0 | 0 |
|  | 24 | 66.2 | ± 1.0 | 34.3 | ± 2.7 | 8.0 | ± 0.3 | 13.0 | ± 0.3 | 0 | 0 |
| Inulin | 2 | 93.7 | ± 0.9 | 30.7 | ± 0.5 | 11.0 | ± 0.0 | 25.0 | ± 0.1 | 20.8 | ± 0.2 |
|  | 4 | 134.0 | ± 2.1 | 52.8 | ± 0.8 | 19.4 | ± 0.6 | 50.8 | ± 1.0 | 0.4 | ± 0.1 |
|  | 8 | 145.1 | ± 1.6 | 58.1 | ± 0.8 | 20.8 | ± 0.2 | 54.2 | ± 0.6 | 0 | 0 |
|  | 24 | 166.8 | ± 3.6 | 71.4 | ± 1.7 | 22.6 | ± 0.5 | 55.2 | ± 1.1 | 0 | 0 |
| Potato starch | 2 | 41.9 | ± 0.3 | 17.6 | ± 0.2 | 7.0 | ± 0.1 | 11.0 | ± 0.1 | 1.0 | ± 0.1 |
|  | 4 | 52.0 | ± 0.1 | 22.7 | ± 0.4 | 9.3 | ± 0.4 | 14.2 | ± 0.4 | 0.7 | ± 0.1 |
|  | 8 | 114.9 | ± 1.6 | 50.3 | ± 0.0 | 23.4 | ± 0.0 | 34.7 | ± 0.4 | 0.7 | ± 0.0 |
|  | 24 | 153.6 | ± 5.0 | 70.7 | ± 0.5 | 26.8 | ± 0.9 | 39.7 | ± 1.6 | 0 | 0 |
| Wheat straw | 2 | 46.0 | ± 0.5 | 21.0 | ± 0.3 | 7.8 | ± 0.0 | 12.0 | ± 0.3 | 0.4 | 0.1 |
|  | 4 | 52.2 | ± 1.8 | 24.7 | ± 0.5 | 8.5 | ± 0.2 | 13.5 | ± 0.4 | 0 | 0 |
|  | 8 | 75.9 | ± 24.0 | 38.2 | ± 0.6 | 11.3 | ± 3.7 | 19.0 | ± 5.9 | 0 | 0 |
|  | 24 | 89.9 | ± 1.2 | 51.5 | ± 2.7 | 9.1 | ± 0.7 | 19.9 | ± 0.2 | 0 | 0 |
| Sugar beet pulp | 2 | 75.9 | ± 1.1 | 36.0 | ± 0.2 | 11.3 | ± 0.6 | 21.3 | ± 0.1 | 1.2 | 0.2 |
|  | 4 | 80.6 | ± 1.3 | 39.7 | ± 0.8 | 11.4 | ± 0.2 | 22.8 | ± 0.4 | 0.2 | 0.0 |
|  | 8 | 106.5 | ± 1.2 | 55.1 | ± 12.2 | 14.3 | ± 0.4 | 29.2 | ± 0.3 | 0 | 0 |
|  | 24 | 159.9 | ± 4.6 | 85.1 | ± 0.7 | 19.2 | ± 1.2 | 40.0 | ± 1.3 | 0 | 0 |

n=2; two-way-ANOVA, R2 adjusted for all tests = 0.99, except for lactic acid. Details of statistical significance are provided in a separate file (see supplementary material file 2: SCFA statistics). The production of lactic acid could not be statistically analyzed due to the lack of measurable values in some of the treatments.

Although statistical analysis could not be conducted due to the absence of lactic acid in some substrates at certain incubation time points, it is noteworthy that after 2 hours of incubation, lactic acid production was observed for inulin, with a value of 20.8 ± 0.2 mg / 200 mg of substrate while other substrates showed almost no lactic acid production during fermentation. Propionic acid levels were significantly increased in inulin compared to the other substrates, peaking at 50.8 mg after 4 hours (*p* < 0.001), and 55.2 mg after 24 hours (*p* < 0.001). Acetic acid was also produced in significant amounts with a peak concentration of 71.4 mg after 24 hours. Straw showed the lowest SCFA concentrations overall, with a total of 89.9 mg at 24 hours (*p* < 0.001). Acetic acid was the most abundant SCFA, reaching 51.5 mg at 24 hours, while butyric acid and propionic acid production was comparatively lower, at 9.1 mg and 19.9 mg respectively. Sugar beet pulp exhibited moderate but steady SCFA production, reaching a total of 159.9 mg at 24 hours. The acetic acid concentration was the highest among all substrates, peaking at 85.1 mg at 24 hours (*p* < 0.001). With 40.0. mg, propionic acid concentration at 24 hours was lower than for inulin (*p* < 0.001), higher than for wheat straw (*p* < 0.001), but did not differ from potato starch (*p* > 0.05). The microbial inoculum alone, without additional substrates, produced the lowest SCFA concentrations, with a total of 66.2 mg at 24 hours (*p* < 0.001).

Acetic acid was the dominant SCFA, with a concentration of 34.3 mg, followed by propionic acid and butyric acid. After 24-hours fermentation, sugar beet pulp showed the highest amount of SCFA, with the highest production of acetic acid (85.1 ml), and high amounts of Butyric acid (19.2 mg) and Propionic acid (40.0).

#### 3.1.3. Bacterial growth based on 16S rRNA genome copies

The total number of bacterial 16S rRNA genome copies was determined using the reference organism *Escherichia coli*, which has seven 16S rRNA gene copies per genome. Figure 3 shows the results of the 16S rRNA genome copy numbers (GCN) of the four different substrates and the inoculum as control over different incubation times (0, 2, 4, 8, 24 hours). Data are given as logarithmic values (log_10_) with Standard deviation (SD) of the measurements, allowing the identification and comparison of differences between the samples. After 2 hours of incubation, the inoculum showed the highest 16S rRNA GCN while potato starch has the second-highest although these differences are not significant (*p* > 0.05, see supplementary material file 3: genome copy numbers). Inulin, wheat straw, and sugar beet pulp exhibited similar GCN values, following comparable trends throughout fermentation. While the inoculum showed a decreasing trend over time (from 8.61 to 8.27, 8.09, and finally 7.21), the other substrates followed a pattern similar to their gas formation. For example, inulin and potato starch reached their peak GCN values after 8 hours, with 8.92 and 8.80 log_10_ GCN, respectively. A similar trend was observed for sugar beet pulp, which peaked at 8 hours with a log_10_ GCN of 8.41.

**Figure 3:**
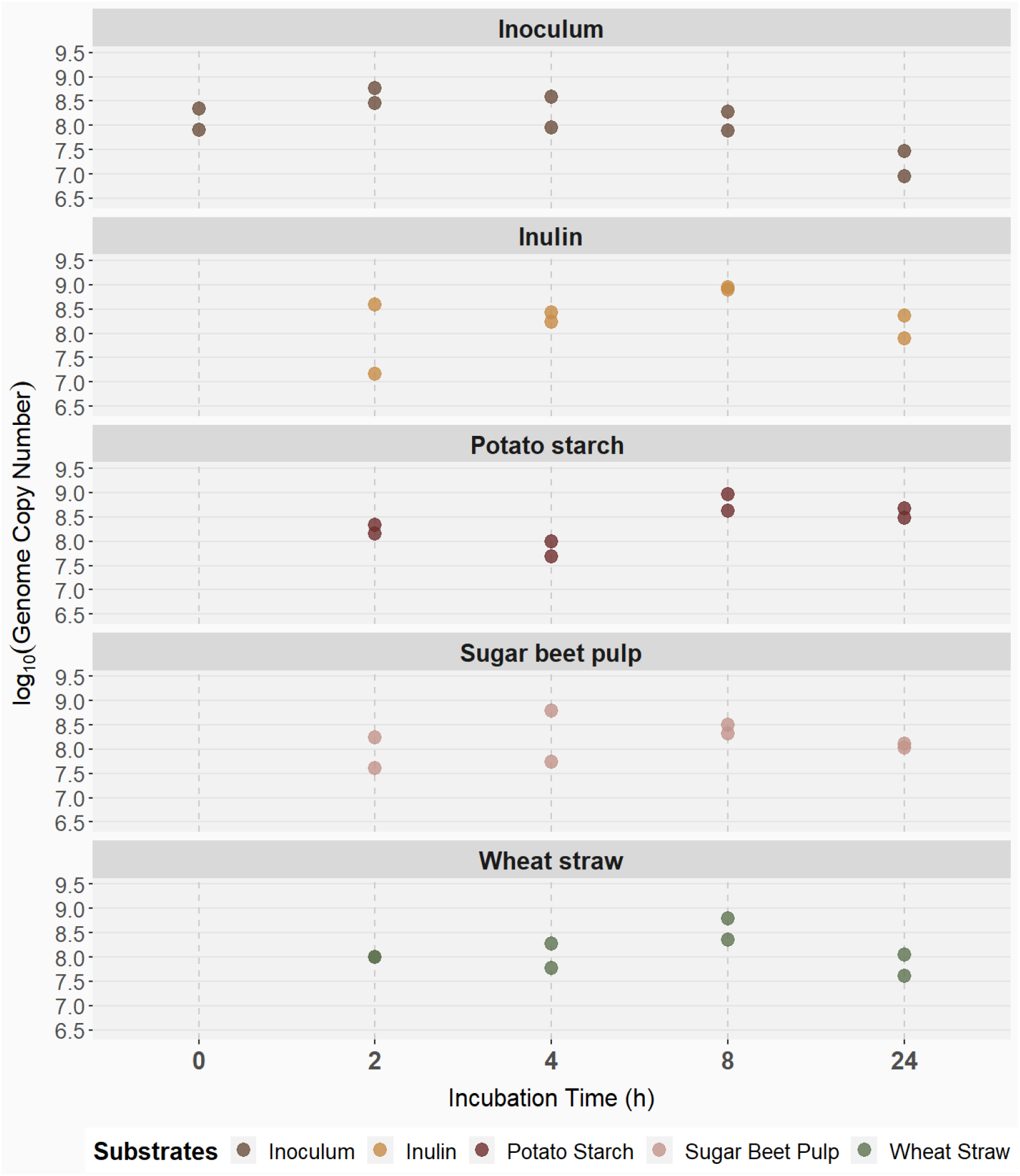
Genome copy numbers (log₁₀) of different substrates (color codes: inulin = yellow; potato starch = dark red; sugar beet pulp = rose; wheat straw = olive) at various incubation times (2, 4, 8, and 24 h). A fecal inoculum from growing pigs was used as a control and analyzed additionally at 0 h. The abscissa represents the incubation time, and the ordinate shows genome copy numbers (log₁₀). Two-way ANOVA was used to assess differences (n = 2). R² = 0.66. No significant differences were observed (*p* > 0.05).

### 3.2. Bacterial community composition and diversity changes in relation to incubated substrates

To assess the bacterial community composition of the different incubation syringes, the individual fecal samples, as well as of the inoculum, 16S rDNA amplicon sequencing of the V3–V4 region was performed. After taxonomic classification, ASVs identified as cyanobacteria or from mitochondrial origins were removed from the dataset. In total, 48 samples were left for further analysis, comprising six individual pig fecal samples, two fecal pool samples mixed with buffer (1:1) and 40 modHGT incubation samples (supernatants of the syringes). The similarity between the inoculum and the microbial profiles of the individual donors served as internal validation, confirming the reliability of the approach and the representativeness of the *in vitro* results (supplementary material file 4: Bacterial composition of the donor pigs)

#### 3.2.1. Alpha diversity

All substrates exhibited comparable bacterial species diversity, with no significant differences in the observed ASVs among the four substrates and the inoculum (*p* > 0.05), as illustrated in the upper part of Figure 4. Differences in alpha diversity were detected in the Shannon index for potato starch after 8 and 24 hours of incubation only, which were significantly different to the other substrates and time points (see supplementary material file 5: Alpha Diversity). Specifically, a decrease in the Shannon index was observed over time for potato starch, with values declining from above 5.5 at 2 and 4 hours to approximately 5 at 8 hours, and further decreasing to below 4.5 at 24 hours of incubation (lower part of Figure 4).

**Figure 4:**
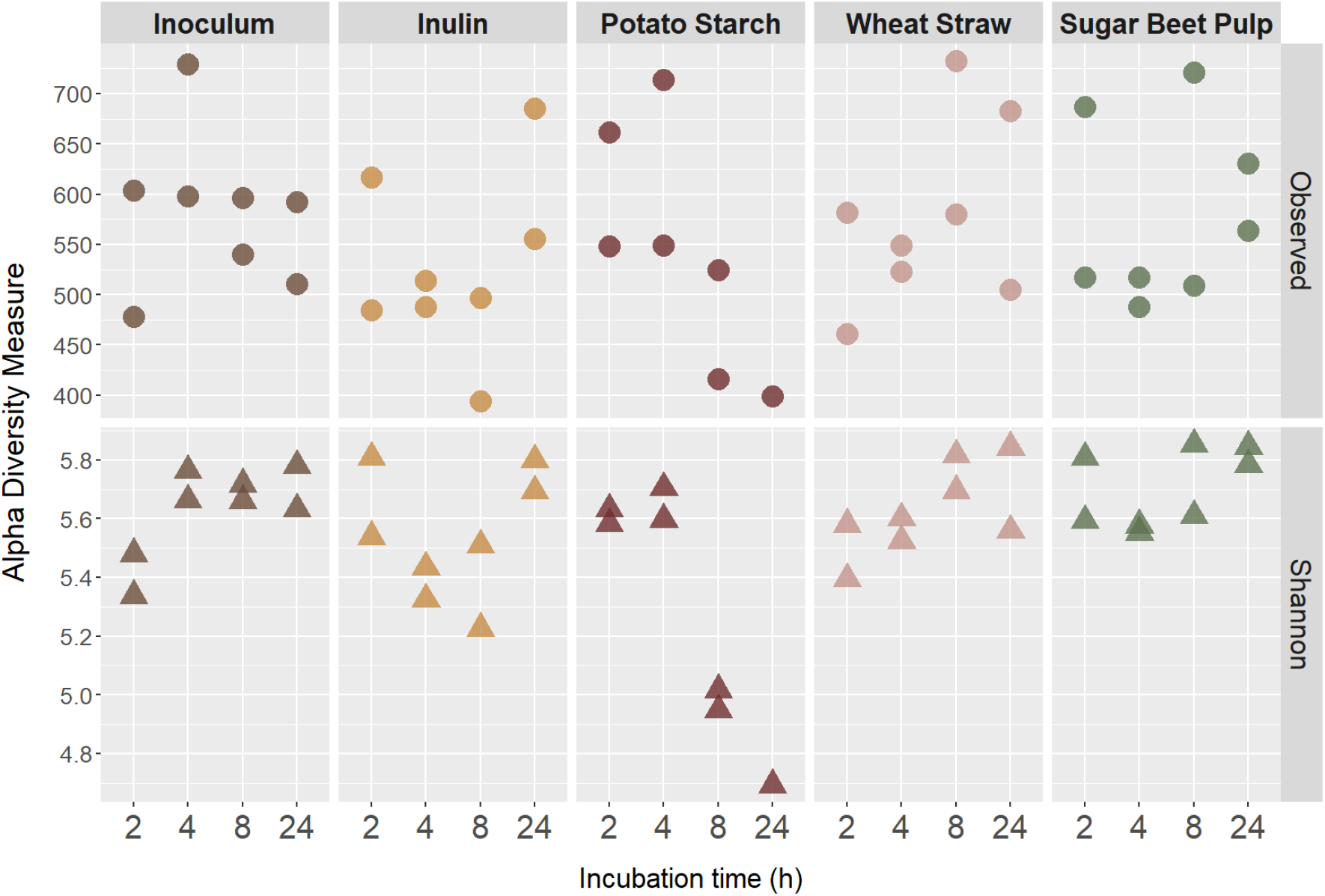
Alpha diversity indices (Observed ASVs and Shannon diversity) of incubation supernatants at different time points (2, 4, 8, 24 h). The abscissa represents the incubation time, and the ordinate shows the respective index values. Color codes: inulin = yellow, potato starch = dark red, wheat straw = rose, sugar beet pulp = olive, fecal inoculum = brown. n = 2, except potato starch at 24 h (n = 1). Results were considered significant at *p* < 0.05. No significant differences were observed in Observed ASVs. Shannon diversity showed significant differences for potato starch at 8 and 24 h (*p* = 0.001).

#### 3.2.2. Beta diversity

Figure 5 illustrates the bacterial community structure over a 24-hour incubation period, assessed using Bray-Curtis dissimilarities on genus level. At each time point, the bacterial communities of the different substrates were significantly distinct. Noticeable changes in bacterial composition were observed as early as 2 hours into incubation. After 4 hours, these shifts became more pronounced, with the substrates diverging further from another. At 8 hours, the differences were even more distinct, and technical replicates of the same substrate clustered closely together. After 24 hours, the inoculum without added substrate remained at the center of the NMDS plot, while the incubated substrates were positioned around it.

**Figure 5:**
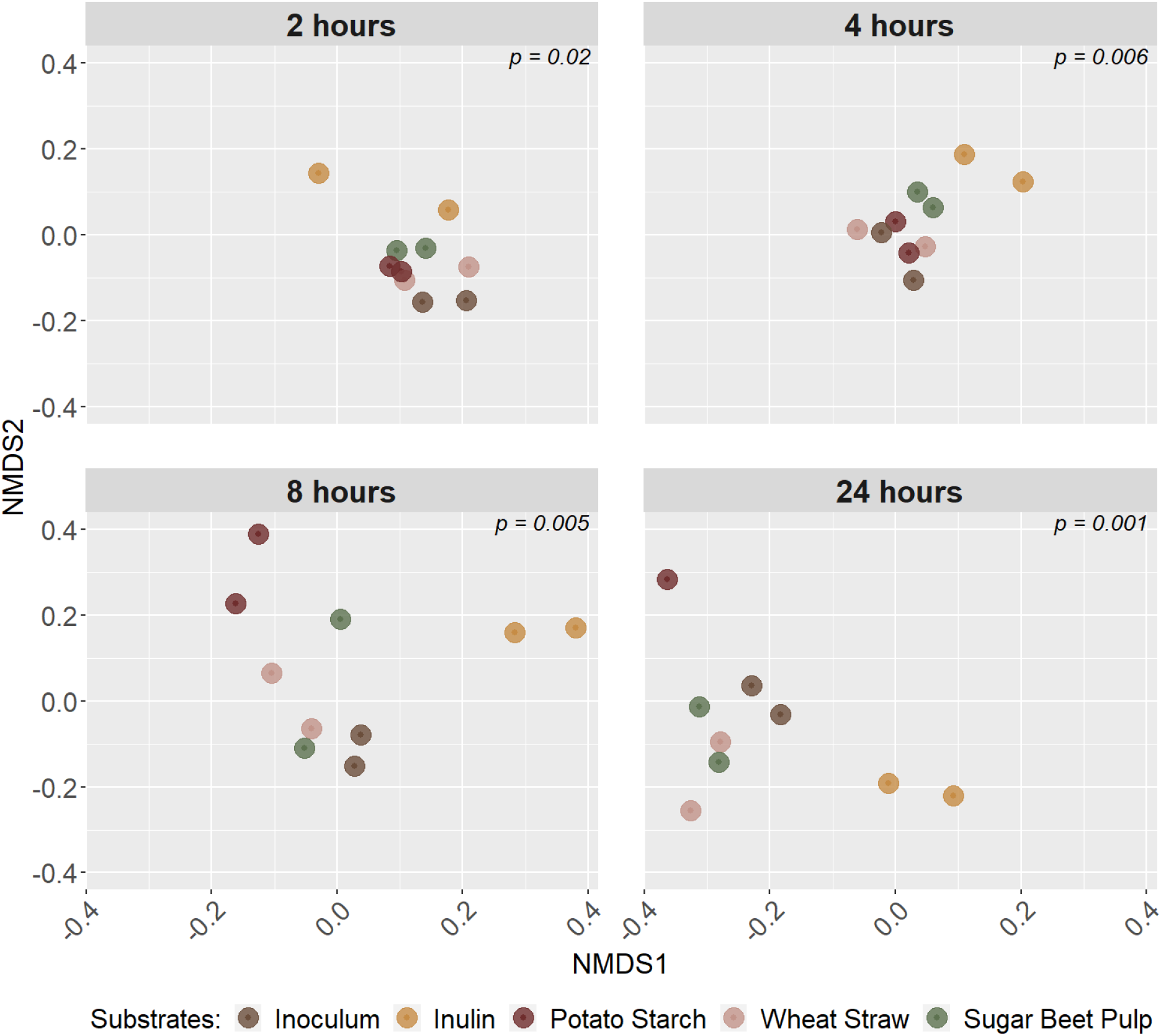
Comparison of microbial community composition (PERMANOVA = Permutational Multivariate Analysis of Variance) across different substrates over 24 h of incubation. NMDS (non-metric multidimensional scaling) plots represent individual time points (2, 4, 8, 24 h) based on genus-level Bray–Curtis dissimilarity. Color codes: inulin = yellow, potato starch = dark red, wheat straw = rose, sugar beet pulp = olive, fecal inoculum = brown. n = 2, except potato starch at 24 h (n = 1). Results were considered significant at *p* < 0.05. Significant differences were already evident at 2 h and persisted throughout (*p* < 0.05).

#### 3.2.3. Taxonomic classification

To gain a deeper understanding of the bacterial community composition at each timepoint of incubation, the differences at the phylum level were examined. Over the course of incubation, regardless of the substrate, Bacteroidota, Firmicutes, and Proteobacteria, Sphirochaetota and Desulfobacteria constituted the three predominant phyla (Figure 6). After 2 and 4 hours of incubation, the samples of all substrates as well as the sample of the inoculum still showed similar proportions of phyla. Differences were observed for potato starch after 8 h of incubation, where Proteobacteria (= red) were over-dominated compared to the other substrates which consisted until the end of incubation (24 h)

**Figure 6:**
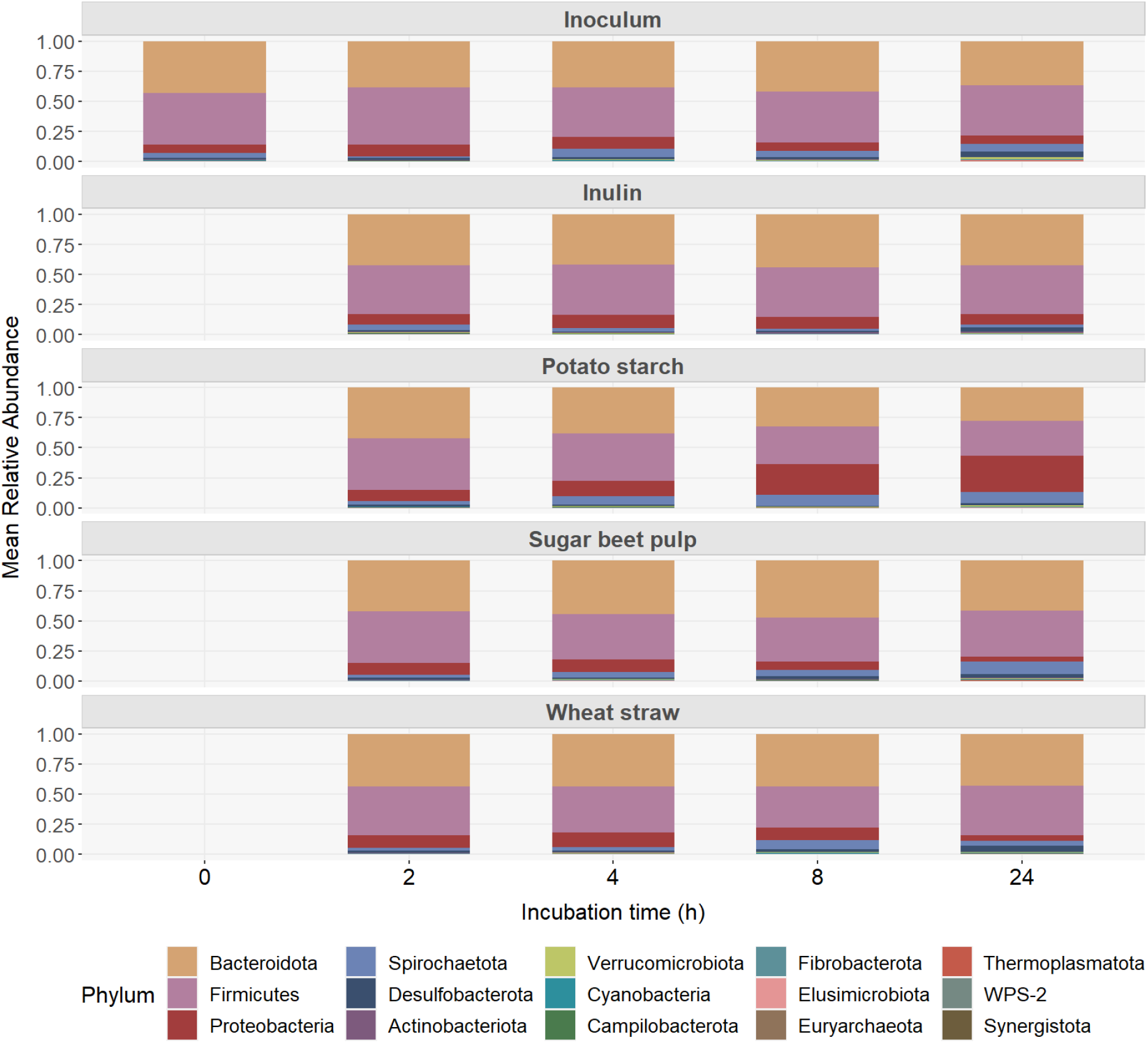
Mean relative abundance of bacterial taxa at phylum level across different substrates (inulin, potato starch, sugar beet pulp, wheat straw) different incubation times (2, 4, 8, and 24 h). The fecal inoculum (internal control) was additionally analyzed at 0 hours. All detected phyla are shown. n = 2, except for potato starch (n = 1).

Furthermore, the bacterial community composition on genus level was examined, where notable variations in community structure were observed (see supplementary material file 6 for detailed relative abundances). Over the 24-hour incubation period, bacterial community composition shifted rapidly in response to the available substrates. At first glance, the incubated substrates exhibited a highly diverse composition, with all genera of the inoculum present in all substrates (Figure 7). Genera with a representation of less than 2% were pooled in “Other”. The starting microbiota composition of all incubated substrates is the inoculum (0-hour time point). After 24 hours of fermentation it included a mix of various genera (158), with noticeable proportions of *Prevotella* (8.96%), *Prevotellaceae_NK3B31_group* (8.10%) *Muribaculaceae (7.10%), Streptococcus* (6.26%) *Alloprevotella* (6.09%), *Rikenellaceae_RC9_gut_group* (4.57%), *Succinivibrionaceae_UCG-001* (4.05%), *Phascolarctobacterium*(3.70%), *Desulfovibrio* (3.64%), *UCG-005* (3.47%), *Prevotellaceae_UCG-003* (3.35%), *Succinivibrio* (3.23%), *Megasphaera* (2.84%), *Treponema* (2.56%), *Lactobacillus* (1.87%) and several other bacteria in lower proportions. After 2 hours of fermentation, the microbiota composition remained relatively close to the original inoculum with some differences, indicating an initial adaptation phase. After 4, 8, and 24 hours of fermentation, there were notable shifts in bacterial composition across all substrates, with certain bacteria becoming more and others less dominant. Overall, one of the most abundant genera was *Muribaculaceae,* which was the predominant genus in the inulin samples after 24 hours of incubation (11.30%) and also highly present in potato starch and wheat straw samples at 24 hours of incubation (7.64 and 7.57%, respectively). In contrast to most genera, that showed a decline in relative abundance over time, the abundance of the genus *Ruminobacter* increased during fermentation. The highest increase was observed for potato starch (0.06% to 25.55%), making it the most dominant genus, followed by Treponema (7.64%) and *Muribaculaceae* (7.57%). For inulin *Escherichia-Shigella* showed a remarkable increase during fermentation. It was initially undetectable, but increased from 0.01% at 8 hours to 2.03% at 24 hours. Furthermore, *Prevotella* and *Megasphaera* showed distinct temporal abundance patterns during fermentation. Both genera increased from 7.96% and 5.85% at 2 hours to 12.81% and 9.44% at 4 hours, respectively, followed by a decline to 9.11% and 0.60% at 24 hours. Wheat straw showed a stable microbial composition after 8 hours, dominated by *Prevotella*, *Prevotellaceae_NK3B31_group*, and *Muribaculaceae*. *Megasphaera* and *Sphaerochaeta* in wheat straw peaked at four to eight hours before decreasing. Predominant after 24 hours were *Rikenellaceae_RC9_gut_group* (11.49%) and *Muribaculaceae* (7.08%) and *Prevotellaceae_NK3B31_group* (6.31%). In sugar beet pulp, *Treponema* increased over the course of fermentation from 1.08% to 7.49% at 24 h. In summary, the microbial composition varied across substrates, with some genera maintaining steady growth while others experienced temporary peaks or declines.

**Figure 7:**
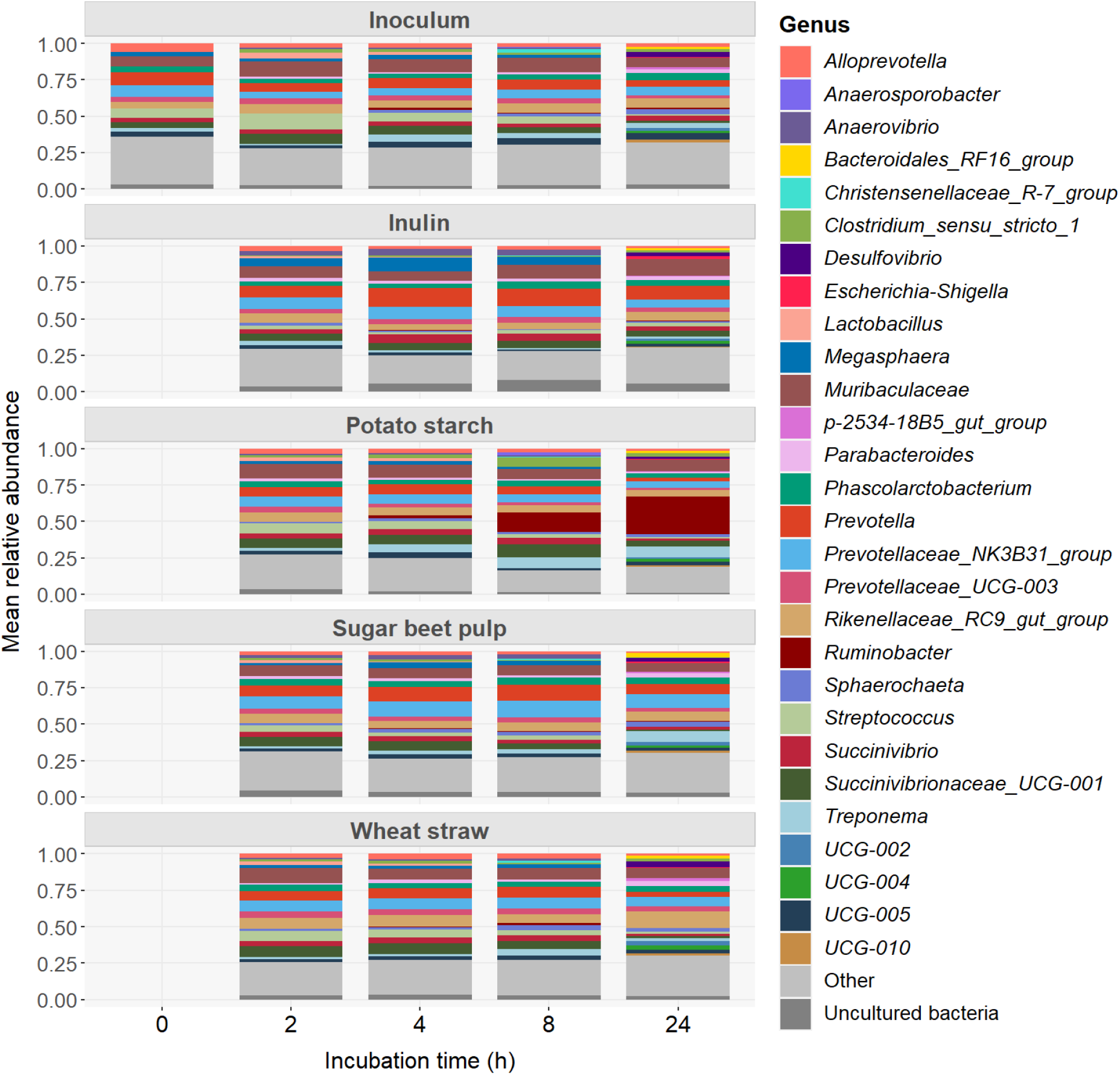
Mean relative abundance of bacterial taxa at genus level across different substrates (inulin, potato starch, sugar beet pulp, wheat straw) at different incubation times (2, 4, 8, and 24 h). The fecal inoculum (internal control) was additionally analyzed at 0 hours. Genera with a relative abundance of < 2% were grouped under “Other”. n = 2, except for potato starch at 24 h (n = 1).

## 4. Discussion

### 4.1. Bacterial community composition and SCFA production

Understanding fermentation processes in the colon of pigs is essential for evaluating the effects of different feeding strategies and substrates. Microorganisms, including bacteria, archaea, and fungi, play a key role in these processes by influencing nutrient digestion and utilisation. In this study, pig feces were used as inoculum for a modHGT to investigate changes in bacterial community composition, gas formation, and SCFA profiles in response to different substrates *in vitro*. The findings contribute to a deeper understanding of pig colon fermentation and its implications for animal nutrition.

Our findings indicate that the bacterial communities and dominant phyla from the incubation experiment are comparable to *in vivo* observations (WYLENSEK ET AL. 2020; KIM AND ISAACSON 2015; LUO ET AL. 2022). However, a closer examination reveals differences in abundances. *In vivo* studies, such as LUO ET AL. (2022), reported that at the phylum level, Firmicutes (68.56%), Bacteroidetes (20.94%), and Spirochaetes (2.59%) were the most abundant. This aligns, in general, with PATIL ET AL. (2020), who described a bacterial community in the colon of healthy adult pigs consisting of 35% Firmicutes, 21% Bacteroidetes, 3% Proteobacteria, and 2% Spirochaetes. Other studies have reported an abundance of Proteobacteria at 13% *(KIM ET AL. 1978; ZHAO ET AL. 2015; PATIL ET AL. 2020)*, which is comparable to our findings, ranging from 4.03% to 12.36%. Overall, our results align with the literature, showing that the three most abundant phyla, regardless of substrate, were Firmicutes (mean: 38.52%), Bacteroidota (mean: 37.47%), and Proteobacteria (mean: 8.20%). These discrepancies may be attributed to variations in fecal sampling methodologies, as rectal swabs primarily capture microbial communities from the distal segment of the gastrointestinal tract. Additional contributing factors include differences in sequencing platforms, the specific hypervariable regions of the 16S rRNA gene targeted (e.g., V1–V2 versus V3–V4), as well as the choice of DNA extraction kits and downstream microbiome profiling pipelines.

In general, we can confirm that the bacterial composition is a typical pig gut community. Microbial activity varied across substrates, as evidenced by gas formation and SCFA concentrations, although the GCN were similar across all substrates and incubation time points. Starting with identical bacterial community across all substrates, the results demonstrated a significantly distinct microbiota already 2 hours after starting the incubation process. This rapid change in the bacterial community depending on the added substrate supports the assumption that available substrates have a strong impact on the microbial community (SCOTT ET AL. 2013). This difference persisted and became more pronounced during incubation, visualized in the NMDS analysis. Even in the inoculum, microbial activity was observed during the 24-hour fermentation period resulting in a notable gas and SCFA production. Nevertheless, bacterial GCN remained unchanged, with only a slight trend noted. Hence, the inoculum, consisting only of buffer and diluted feces, contained sufficient residual nutrients to sustain microbial activity without additional substrate. Some bacteria increased (i.e., *Rikenellaceae_RC9_gut_group*) while other decreased (i.e., *Streptococcus)* over the course of fermentation. *Streptococcus* is known for glucose fermentation resulting in lactic acid as end product *(DU TOIT ET AL. 2014)* explaining higher concentrations of lactic acid in inoculum after 2 hours of fermentation. As the proportion of *Streptococcus* decreased, lactic acid levels declined towards zero during incubation, likely because it was utilised as a substrate by other bacteria.

Compared to the other substrates, *Megasphaera* was one of the dominant genera that was formed during the incubation of inulin (5.85% after 2 hours of incubation). *Megasphaera elsdenii* is a lactate-utilizing bacterium commonly found in the intestines of pigs, where it is colon-associated, and is capable of producing SCFA such as acetic acid, propionic and butyric acid (KAJIHARA ET AL. 2017; YOSHIKAWA ET AL. 2018; HOLMAN ET AL. 2017). In a study by YOSHIKAWA ET AL. (2018), the Y2 strain of *Megasphaera elsdenii* rapidly utilised the complete lactate thereby producing acetate and propionate during 6 hours of incubation. In this study, during the first 4 hours of the fermentation of inulin, the total SCFA concentration and in particular the concentration of acetic acid and propionic acid increased, in line with the abundance of *Megasphaera*. However, after reaching a peak abundance of 9.44% at 4 hours, the amount of *Megasphaera* decreased to only 0.6% after 24 hours being outcompeted by other bacterial taxa in the later stages of incubation. This suggests that the presence of this bacterium is dependent on the availability of lactic acid, which is only available for a short period of time due to the batch approach of the modHGT and therefore may not accurately reflect the situation *in vivo*. Compared to the other substrates, inulin exhibited a higher lactic acid concentration after 2 hours of incubation (20.85 mg / 200 mg DM) thereby favouring the growth of *Megasphaera elsdenii* and possibly impairing that of other microbial genera due to the conversion of lactic acid into other SCFA and the accompanied decrease in pH. The noticeable lactic acid production during early incubation of inulin under the conditions of the modHGT indicates a positive impact on the growth of lactic acid producing bacteria, which may warrant further investigation. Lactic acid producing bacteria are known for improving gut health by regulating the intestinal environment, inhibiting pathogens, enhancing microbial balance, and supporting immune function (YANG ET AL. 2015; PESSIONE 2012). After 8 hours of fermentation, compared to the other treatments, the most unclassified genera were observed in the inulin samples and might indicate unique bacterial response to inulin and further potential for investigation of these unclassified bacteria. *Phascolarctobacterium* found in fermented inulin, is a genus of bacteria, including *Phascolarctobacterium succinatutens*, which is a strictly anaerobic bacterium that uses succinate as its main carbon source and can produce SCFA, including acetic acid and propionic acid (WU ET AL. 2017; WATANABE ET AL. 2012). It may play a role in the gut microbiome and is cecum associated (HOLMAN ET AL. 2017), although its precise physiological functions require further investigation (CRESPO-PIAZUELO ET AL. 2019).

In monogastric animals, symbiotic digestion occurs through postgastric fermentation. Food is first acidically digested in the stomach, where hydrolysis products, including soluble sugars, are absorbed. Undigested plant polymers are then fermented in an elongated colon or cecum (or both) where they are converted to SCFA such as acetic acid, propionic acid and butyric acid. These fatty acids are absorbed by the host animal and serve as an important source of energy. Postgastric fermentation is the predominant fermentation strategy in mammals (Bacterial Biogeochemistry 2012).

*In vivo*, **potato starch** is digested in the small intestine. The rapid increase in acetate production observed between 8 and 24 hours of fermentation in potato starch may be explained by the substantial proliferation of *Ruminobacter*. This obligate anaerobe uses only α-linked glucose molecules, present like in starch, as a source of energy and is known for producing mainly formate, acetate and succinate as end products and thus excellent model for the study of bacterial starch degradation (ANDERSON 1995). Analysis of the abundance plot reveals that almost all microbial genera experienced a reduction in abundance after 8 hours of fermentation. This decline is likely attributed to the extreme dominance of *Ruminobacter*, whose abundance increased from 0.06% after 2 hours to 2.38% after 4 hours, 13.48% after 8 hours, and ultimately 25.55% after 24 hours of incubation. To further investigate what consequences dominance of *Ruminobacter* has on the gut microbiota and metabolites, it is advisable to predigest starch and then ferment it in the modHGT to analyse microbial degredation of starch. The highest gas formation was observed with potato starch after 24 hours of fermentation. During the initial stages of the fermentation process, acetic acid levels were relatively low compared to other substrates. However, a significant increase in acetic acid concentration was observed after 8 hours, probably due to the activity of *Ruminobacter,* which showed a marked increase in abundance during this period. *Ruminobacter* are known to produce SCFA, including acetic acid (HUA ET AL. 2022).

**Wheat straw**, known for its complex carbohydrate structure, consisting of x, y, and z (ZIKELI ET AL. 2016), may require specialised degraders and act more slowly (LKHAGVA ET AL. 2021), resulting in less gas formation and less SCFA production due to delayed microbial activity, even when there was no significant difference in GCN or observed genera compared to the other substrates. The bacterial community in fermented wheat straw was more stable over the course of fermentation with regard to the changes in abundance of the bacteria. High fiber levels are known to stimulate the growth of cellulolytic and xylanolytic bacteria (Moore et al., 1987; Durmic et al., 1998). As a lignocellulosic substrate, wheat straw supports the growth of bacterial genera capable of degrading fibrous materials, such as Rikenellaceae_RC9_gut_group and Christensenellaceae. *Succinivibrionaceae_UCG-001* decreased from 8 to 24 h, *Rikenellaceae_RC9_gut_group, Desulfovibrio* increased. High levels of fiber have been found to stimulate the growth of genera with cellulolytic and xylanolytic activities (MOORE ET AL. 1987; DURMIC ET AL. 1998). Consistent with that the delayed and lower total SCFA production in wheat straw fermentation suggests a less efficient fermentation process compared to other substrates. Notably, SCFA composition changed primarily between 8 and 24 hours, with an increase in acetic acid, while propionic and butyric acid levels remained largely unchanged. This indicates that acetate-producing bacteria were dominant. One notable genus, *Treponema*, showed a peak abundance of 4.58% after 8 hours, decreasing to 1.82% at 24 hours. In the same time, acetic acid was produced probably because of *Treponema*. Treponema has an important role in fiber digestion and energy supply to the host (GARDINER ET AL. 2020).

Beta-diversity of fermented **sugar peet pulp** showed most similarities with wheat straw, but was still significantly different to it. Several bacteria, including *Christensenellaceae* and *Treponema*, contribute to the breakdown of plant polysaccharides to produce short-chain fatty acids (SCFAs) as an energy source for pigs, while *Treponema* improve fiber digestibility by converting indigestible compounds into metabolisable energy (GARDINER ET AL. 2020). In fermented sugar beet pulp, *Treponema* had an abundance of almost 10% after 24 hours of fermentation. There is evidence, that a diet containing low proportions (3%) of sugar beet pulp could be beneficial for the pig’s gut health but more research is needed to investigate, if there are antinutritive factors with a negative impact (BADARAS ET AL. 2022). Several microorganisms, such as different cellulose degraders, are functionally redundant (Bacterial Biogeochemistry 2012). Therefore, it is plausible that especially in fermented sugar beet pulp and wheat straw, similar genera act as fiber degraders, resulting in the most similar bacterial community in the NMDS plot over the course of fermentation, while still being significantly different. In general, the bacterial composition of the substrates was highly diverse, with many bacteria coexisting in what could be described as a highly diverse bacterial ecosystem. Compositional analysis further confirmed that different fiber types and carbohydrates selectively enrich specific bacterial genera, consistent with observations from *in vivo* studies (UMU ET AL. 2015). During fermentation, the composition of the microbiota shifts as it dynamically adapts to each substrate, with some genera showing a clear preference for a particular substrate as indicated by relative abundance plots. These changes in substrate availability induced a rapid bacterial response, influencing both microbial abundance and metabolite production such as SCFAs. Different bacterial taxa exhibited distinct growth patterns, with some peaking at 8 hours before declining, while others reached their highest abundance after 24 hours. This suggests potential metabolic interdependencies, where certain bacteria utilize metabolites produced by earlier-peaking species, thereby promoting subsequent microbial growth.

### 4.2. Challenges and applicability of the study

One of the major challenges of modHGT is the necessity for an incubation environment that closely resembles *in vivo* conditions. Organisms unable to grow under these conditions are not represented (SCOW ET AL. 2001) and certain bacterial genera, that are closely associated with the gut mucosa are excluded due to the sampling method and, as a result, not present in the fecal inoculum. In the context of a healthy microbial community, which is characterized by diversity, stability, and resilience to minor changes (LEVY ET AL. 2017), it appears that the bacterial composition in fecal samples lacks such stability and resilience. This is evidenced by observable changes occurring within the first 2 hours of fermentation. However, it is plausible that certain bacterial communities exhibit greater resistance to changes, potentially due to the presence of a constant factor for example the mucosal environment. This stable parameter may provide a consistent niche for specific bacteria, contributing to their relative stability despite external variations like the available substrates (nutrients). It seems that the modHGT is much more sensitive to changes than the *in vivo* system. It is important to note that the *in vitro* batch culture system, like the modHGT, represents a static rather than a dynamic environment. *In vivo*, certain nutrients, particularly SCFAs, would be absorbed by the intestinal epithelium or buffered within the lumen to maintain a relatively stable pH. In contrast, such regulatory mechanisms are absent in the glass syringes, leading to the accumulation of SCFAs over time. As a result, pH may shift despite the presence of a buffer solution, potentially affecting the growth and survival of specific bacterial taxa.

Addressing these limitations is essential for improving the accuracy and applicability of modHGT results. A promising aspect for future research is investigating gut microbiota development across different growth stages in pigs. It is well established that microbial diversity changes significantly from birth to post-weaning (INOUE ET AL. 2005). While breed has a relatively minor impact on gut bacterial succession (BIAN ET AL. 2016), it remains an important factor that could be explored further. By studying these variations, researchers can gain deeper insights into microbiota development throughout a pig’s lifespan and production cycle and its implications for different feeding strategies and therefore health and productivity. In addition to its potential for fundamental microbiota research, the modHGT system offers promising applicability as a pre-screening tool for microbial or nutritional interventions, such as probiotics, prebiotics, or dietary additives. Due to its reproducibility, controlled conditions, and substrate-specific responsiveness, the system allows for a preliminary assessment of microbial shifts and fermentation dynamics prior to conducting *in vivo* trials. This can reduce experimental costs, and the number of animals used, aligning with the principles of the 3Rs (Replacement, Reduction, Refinement). Especially in early development stages, the modHGT facilitates the identification of promising approaches for further validation.

The microbial communities in the gastro intestinal tract are influenced by the host animal’s diet, and this, in turn, affects the distribution of microbiota within the gastro intestinal tract (LESER ET AL. 2002; DORÉ AND BLOTTIÈRE 2015). During incubation, this microbial adaptation was reflected in the differential responses of bacterial species to substrates (their diet). The relative abundance of certain bacterial genera along the gastrointestinal tract is likely to be influenced by the physiological conditions in each section. For example, certain bacterial species prefer certain pH values, oxygen levels and nutrient sources, which may differ between sections of the digestive tract (LKHAGVA ET AL. 2021). If only one substrate is present, as in the modHGT, this can influence the composition of the bacterial community, as bacteria can adapt and multiply to use this substrate efficiently. A substrate-specific enrichment of bacterial genera is evident, as each substrate promotes the growth of distinct bacterial groups. Certain dominant species, particulary among the Bacteroidota, are known to switch between different energy sources in the gut depending on availability (FLINT ET AL. 2012). For example, *Prevotella* and *Anaerosporobacte*r appear to be generalists that can grow on multiple substrates. In contrast, some genera that can process the substrate most efficiently, such as *Ruminobacter*, are overrepresented on rapidly fermentable, glucose-containing substrates such as potato starch (Figure 7).

Even when a particular substrate is dominant, the bacterial consortium can interact with each other either through direct competition for ecological niches and nutrients, leading to the dominance or subordination of a few genera, or through indirect mechanisms, such as the release of metabolic products that affect other bacteria (LOUIS ET AL. 2007). In the colon, certain bacterial species are examples of these interactions, surviving by cross-feeding where they use either the breakdown products of complex carbohydrates or fermentation products such as lactic acid to support their growth (SCOTT ET AL. 2013). But compared to *in vivo*, there is no interaction between the bacteria and their host in the modHGT. The limitations of batch systems such as the HGT for the simulation of rumen conditions are well documented in the literature, but it is possible to transfer these findings to the modHGT using pig feces. These systems lack the dynamic interactions that occur in a living host, such as nutrient absorption, mineral exchange and excretion of waste metabolites. This leads to an accumulation of metabolites and changes in the substrate environment over time which can affect microbial activity and fermentation outcomes, such as gas production or SCFAs (WISCHER ET AL. 2013a).

Looking at the starter culture for the modHGT, both the pool-feces-buffer mixture and the inoculum were effective in representing the diversity found in the individual donor pig samples. This positioning suggests that while the inoculum is not precisely centered among the individual fecal samples, it reflects a different but related bacterial community structure. Technical replicates of the inoculum clustered closely together, underscoring the consistency and reproducibility of the analytical process well as the method of the modHGT. This variability reflects the natural diversity among individual pig samples. The pool-feces-buffer mixture was representative of the diversity found in the individual samples, effectively capturing the range of observed genera.

## 5. Conclusion and outlook

Regarding the 3R principals and the reduction of animal studies, a promising approach for microbiome research would be the identification and assembly of an ‘ideal *in vivo*-like’ microbiome *in vitro*. By cultivating representative strains of porcine gut microbiota without the constant need for samples from live animal, researchers could establish a stable microbial community for incubation studies. This requires a comprehensive understanding of the bacterial genera present in the gut. The modHGT could provide insights in functional aspects of gut microbes and perhaps host interactions within such a standardized microbiome. A comparable approach has already been implemented in human gut microbiota research, where standardized microbiome reference communities, such as the ZymoBIOMICS Gut Microbiome Standard (Zymo Research, Irvine, CA, USA), are used. Applying similar standardization to pig microbiota studies could enhance comparative research while accounting for breed-specific, dietary, and environmental variations. For instance, when studying specific breeds like the Angler Saddleback pig, microbiome standards could be adapted to address targeted research questions more effectively and across different farm settings that change microbiota, breed-specific differences allowing for more precise comparative research.

The modHGT is a valuable tool to improve our understanding of the porcine gut microbiota. This method is accessible, requires minimal input, and aligns with animal welfare considerations. It holds great potential for standardising microbiota research and could contribute to refining knowledge of feeding strategies for pigs (ZHANG ET AL. 2024). By utilizing the modHGT, researchers could develop targeted feed formulations and optimise farming practices to improve efficiency and sustainability. It can also help refine the knowledge graph of pig gut microbiota and pig feeding of ZHANG ET AL. (2024). This is particularly relevant in the context of regulatory changes, such as the ban on veterinary drugs containing high doses of zinc oxide (3000 mg/kg) by the European Comission in 2022. These restrictions, aiming to reduce antimicrobial resistance and environmental contamination, require alternative strategies for maintaining the health and growth of pigs. *In vitro* tools like the modHGT help to gain insights into the development of the gut microbiota under different feeding strategies and could offer innovative and sustainable solutions to meet these challenges. Ultimately, the gut microbiota and its host form a complex, interdependent system with archaea and fungi also play a role in the fermentation and digestion processes, which were not further considered in this study but also contribute to the gastrointestinal tract ecosystem. Future research should investigate the consequences of altering the bacterial composition within this system and its physiological functions and consequences.

## Acknowledgement

The authors wish to thank the ISF Schaumann Forschung GmbH (Wahlstedt, Germany) for the conceptualization of this study, the HPLC analysis of the SCFA and for financial support in sequencing the microbiome data. The authors would like to thank Mario Hasler (CAU) for his advice regarding the statistical analysis as well as the staff of the microbiome lab of the Institute of Clinical Molecular Biology, for their assistance during laboratory procedures. Microbiota sequencing received infrastructure support from the DFG Excellence Cluster 2167 “Precision Medicine in Chronic Inflammation” (PMI) and the DFG Research Unit 5042 „miTarget”.

## Declaration of generative AI and AI-assisted technologies in the writing process

During the preparation of this work the authors used *deepl.write* and *chatGPT* for English language editing. After using these tools, the authors reviewed and edited the content as needed and take full responsibility for the content of the published article.

