## Supplementary File 5 for "From Substrate to Short Chain Fatty Acids: An *In Vitro* Investigation of Bacterial Dynamics in the Porcine Colon using the Modified Hohenheim Gas Test"

**Table S18:** Summary of shannon index values during *in vitro* fermentation of different substrates using the modHGT with pig feces at 2, 4, 8, and 24 hours of incubation (n = 2; potato starch 24h: n = 1)

| Substrate | 2h | | 4h | | 8h | | 24h | |
|---|---|---|---|---|---|---|---|---|
| Inoculum | 5.41 | ± 0.1 | 5.71 | ± 0.07 | 5.69 | ± 0.04 | 5.71 | ± 0.11 |
| Inulin | 5.67 | ± 0.19 | 5.38 | ± 0.08 | 5.37 | ± 0.20 | 5.75 | ± 0.08 |
| Potato starch | 5.61 | ± 0.04 | 5.65 | ± 0.08 | 4.98 | ± 0.05 | 4.69 | na |
| Wheat straw | 5.49 | ± 0.13 | 5.56 | ± 0.5 | 5.76 | ± 0.09 | 5.70 | ± 0.20 |
| Sugar beet pulp | 5.7 | ± 0.15 | 5.57 | ± 0.02 | 5.73 | ± 0.17 | 5.81 | ± 0.04 |

Mean values ± standard deviation are shown.

**Table S1:** Statistical comparisons of Shannon index values between substrates during *in vitro* fermentation (general linear hypothesis).

| Linear Hypotheses | Estimate | se | t value | Pr(>\|t\|) | |
|---|---|---|---|---|---|
| inoculum - inulin, 2 == 0 | -0.263915 | 0.114161 | -2.312 | 0.4957 | |
| inoculum - starch, 2 == 0 | -0.197723 | 0.114161 | -1.732 | 0.8523 | |
| inoculum - straw, 2 == 0 | -0.078493 | 0.114161 | -0.688 | 10.000 | |
| inoculum - sugar beet pulp,2 == 0 | -0.291793 | 0.114161 | -2.556 | 0.3518 | |
| inulin - starch, 2 == 0 | 0.066192 | 0.114161 | 0.580 | 10.000 | |
| inulin - straw, 2 == 0 | 0.185422 | 0.114161 | 1.624 | 0.8996 | |
| inulin - sugar beet pulp, 2== 0 | -0.027878 | 0.114161 | -0.244 | 10.000 | |
| starch - straw, 2 == 0 | 0.119230 | 0.114161 | 1.044 | 0.9976 | |
| starch - sugar beet pulp, 2== 0 | -0.094070 | 0.114161 | -0.824 | 0.9998 | |
| straw - sugar beet pulp, 2== 0 | -0.213300 | 0.114161 | -1.868 | 0.7792 | |
| inoculum - inulin, 24 == 0 | -0.040868 | 0.114161 | -0.358 | 10.000 | |
| inoculum - starch, 24 == 0 | 1013693 | 0.139818 | 7250 | <0.001 | *** |
| inoculum - straw, 24 == 0 | 0.004964 | 0.114161 | 0.043 | 10.000 | |
| inoculum - sugar beet pulp,24 == 0 | -0.108671 | 0.114161 | -0.952 | 0.9991 | |
| inulin - starch, 24 == 0 | 1054561 | 0.139818 | 7542 | <0.001 | *** |
| inulin - straw, 24 == 0 | 0.045832 | 0.114161 | 0.401 | 10.000 | |
| inulin - sugar beet pulp, 24 == 0 | -0.067803 | 0.114161 | -0.594 | 10.000 | |
| starch - straw, 24 == 0 | -1008729 | 0.139818 | -7215 | <0.001 | *** |
| starch - sugar beet pulp, 24 == 0 | -1122364 | 0.139818 | -8027 | <0.001 | *** |
| straw - sugar beet pulp, 24== 0 | -0.113635 | 0.114161 | -0.995 | 0.9986 | |
| inoculum - inulin, 4 == 0 | 0.334366 | 0.114161 | 2.929 | 0.1892 | |
| inoculum - starch, 4 == 0 | 0.061937 | 0.114161 | 0.543 | 10.000 | |
| inoculum - straw, 4 == 0 | 0.150276 | 0.114161 | 1.316 | 0.9788 | |
| inoculum - sugar beet pulp,4 == 0 | 0.147694 | 0.114161 | 1.294 | 0.9817 | |
| inulin - starch, 4 == 0 | -0.272429 | 0.114161 | -2.386 | 0.4489 | |
| inulin - straw, 4 == 0 | -0.184091 | 0.114161 | -1.613 | 0.9040 | |
| inulin - sugar beet pulp, 4== 0 | -0.186672 | 0.114161 | -1.635 | 0.8952 | |
| starch - straw, 4 == 0 | 0.088338 | 0.114161 | 0.774 | 0.9999 | |
| starch - sugar beet pulp, 4== 0 | 0.085757 | 0.114161 | 0.751 | 0.9999 | |
| straw - sugar beet pulp, 4== 0 | -0.002582 | 0.114161 | -0.023 | 10.000 | |
| inoculum - inulin, 8 == 0 | 0.322138 | 0.114161 | 2.822 | 0.2288 | |
| inoculum - starch, 8 == 0 | 0.707081 | 0.114161 | 6194 | <0.001 | *** |
| inoculum - straw, 8 == 0 | -0.066420 | 0.114161 | -0.582 | 10.000 | |
| inoculum - sugar beet pulp,8 == 0 | -0.042134 | 0.114161 | -0.369 | 10.000 | |
| inulin - starch, 8 == 0 | 0.384943 | 0.114161 | 3.372 | 0.0819 | . |
| inulin - straw, 8 == 0 | -0.388559 | 0.114161 | -3.404 | 0.0770 | . |
| inulin - sugar beet pulp, 8== 0 | -0.364272 | 0.114161 | -3.191 | 0.1167 | |
| starch - straw, 8 == 0 | -0.773502 | 0.114161 | -6776 | <0.001 | *** |
| starch - sugar beet pulp, 8== 0 | -0.749215 | 0.114161 | -6563 | <0.001 | *** |
| straw - sugar beet pulp, 8== 0 | 0.024287 | 0.114161 | 0.213 | 10.000 | |

Two-way ANOVA was performed with substrate and incubation time as factors. Adjusted p-values (single-step method) are reported. Adjusted $R^2 = 0.90$. Comparison of substrate levels included. se = standard error of the estimate, Significance codes: *** $p < 0.001$, ** $p < 0.01$, * $p < 0.05$, · $p < 0.1$
