## Supplementary File 4 for "From Substrate to Short Chain Fatty Acids: An *In Vitro* Investigation of Bacterial Dynamics in the Porcine Colon using the Modified Hohenheim Gas Test"

Six individual fecal samples were collected from the donor pigs. Additionally, replicates of both the pool-feces-buffer mixture (dilution 1:1) and the inoculum used for incubation were sequenced. The Alpha diversity of individual fecal samples from the donor pigs varied significantly $p < 0.05$, with observed genera ranging from approximately 350 to over 600. While the inoculum displayed a higher count of observed genera relative to the individual fecal samples, this difference was not statistically significant ($p > 0.05$).

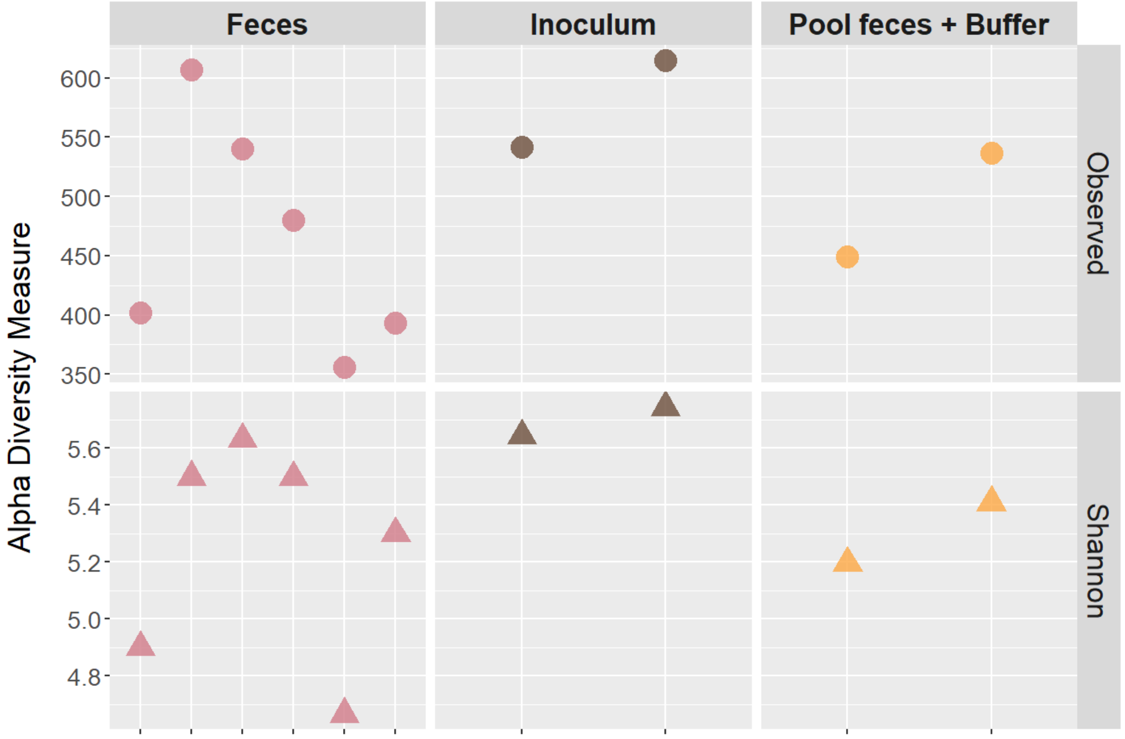


**Figure S1:** Alpha diversity indices (observed ASVs and Shannon diversity) of individual feces, Inoculum used for incubation and pool feces – buffer with 1:1 dilution, with the abscissa representing incubation time (0) and the ordinate indicating the respective indices. Individual feces n=6; Inoculum and Pool Feces + Buffer n=2.

The genus-level microbiota composition varied significantly among the individual fecal samples from donor pigs (Figure S2**Fehler! Verweisquelle konnte nicht gefunden werden.**). As expected, the individual fecal samples were dispersed around the inoculum and the pooled feces-buffer mixture in the ordination plots.

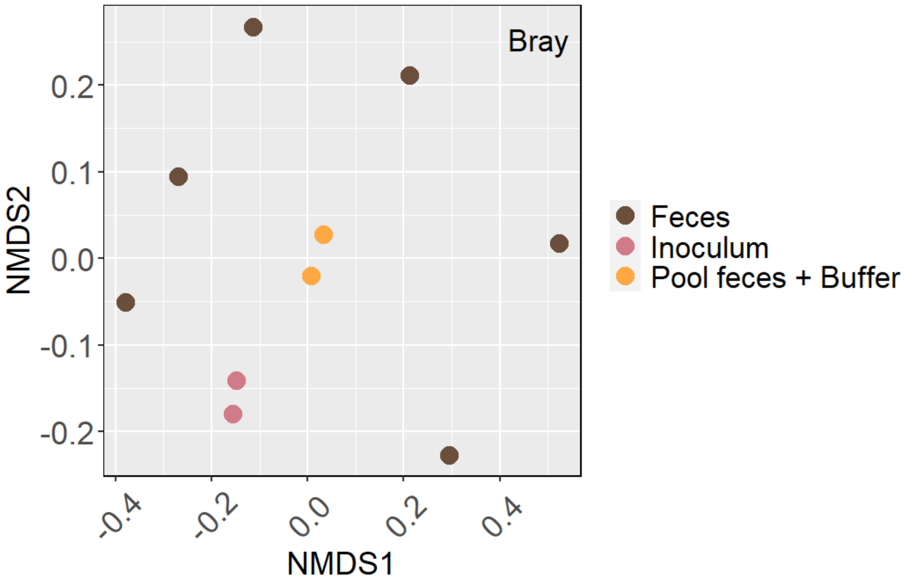


**Figure S2**: Comparison of microbial community composition (PERMANOVA) of individual fecal samples, pooled feces–buffer mixture, and inoculum at 0 hours, based on genus-level Bray–Curtis distances visualized by NMDS (Non-metric Multidimensional Scaling). Individual feces = brown (n = 6); inoculum = pink (n = 2); pooled feces + buffer = orange (n = 2). Significant differences were considered at $p < 0.05$. No significant differences were found.
