## Supplementary File 3 for "From Substrate to Short Chain Fatty Acids: An *In Vitro* Investigation of Bacterial Dynamics in the Porcine Colon using the Modified Hohenheim Gas Test"

**Table S7:** Summary of 16S rRNA genome copy numbers ($log_{10}$) during *in vitro* fermentation of different substrates using the modHGT with pig feces at 2, 4, 8, and 24 hours of incubation. Values were calculated assuming seven 16S rRNA gene copies per bacterial genome (based on *E. coli*).

| Substrate n = 2 | t (h) | 16S rRNA genome copies [log10] | ± SD |
|---|---|---|---|
| Potato starch | 2 | 8.25 | ± 0.12 |
| | 4 | 7.84 | ± 0.22 |
| | 8 | 8.80 | ± 0.24 |
| | 24 | 8.58 | ± 0.14 |
| Inulin | 2 | 7.88 | ± 1.00 |
| | 4 | 8.33 | ± 0.14 |
| | 8 | 8.92 | ± 0.03 |
| | 24 | 8.13 | ± 0.33 |
| What Straw | 2 | 8.00 | ± 0.01 |
| | 4 | 8.02 | ± 0.36 |
| | 8 | 8.58 | ± 0.30 |
| | 24 | 7.83 | ± 0.31 |
| Sugar beet pulp | 2 | 7.93 | ± 0.44 |
| | 4 | 8.26 | ± 0.74 |
| | 8 | 8.41 | ± 0.13 |
| | 24 | 8.07 | ± 0.06 |
| Inoculum | 2 | 8.61 | ± 0.23 |
| | 4 | 8.27 | ± 0.45 |
| | 8 | 8.09 | ± 0.28 |
| | 24 | 7.21 | ± 0.37 |

Mean values ± standard deviation (SD) are shown; t = Incubation time

**Comparison of incubation times:** To investigate temporal dynamics in bacterial genome copy numbers, estimated marginal means (EMMs) for incubation time were computed, stratified by substrate. A linear model was fitted to $\log_{10}$-transformed genome copy numbers, including incubation time, substrate, and their interaction as fixed effects. Pairwise comparisons between incubation time points were conducted to identify statistically significant differences. No significant differences were found.

Table S8: Estimated Marginal Means ($\log_{10}$) of 16S rRNA Genome copy numbers over time during *in vitro* fermentation of the **inoculum**. Pairwise comparisons between incubation times.

| contrast | estimate | se | df | t.ratio | p.value |
|---|---|---|---|---|---|
| incub_time0 - incub_time2 | -0.4847 | 0.404 | 27 | -1.199 | 0.7521 |
| incub_time0 - incub_time4 | -0.1396 | 0.404 | 27 | -0.345 | 0.9968 |
| incub_time0 - incub_time8 | 0.0439 | 0.404 | 27 | 0.109 | 1.0000 |
| incub_time0 - incub_time24 | 0.9243 | 0.404 | 27 | 2.286 | 0.1804 |
| incub_time2 - incub_time4 | 0.3450 | 0.404 | 27 | 0.853 | 0.9112 |
| incub_time2 - incub_time8 | 0.5286 | 0.404 | 27 | 1.307 | 0.6891 |
| incub_time2 - incub_time24 | 1.4090 | 0.404 | 27 | 3.484 | 0.0135 |
| incub_time4 - incub_time8 | 0.1836 | 0.404 | 27 | 0.454 | 0.9907 |
| incub_time4 - incub_time24 | 1.0639 | 0.404 | 27 | 2.631 | 0.0926 |
| incub_time8 - incub_time24 | 0.8804 | 0.404 | 27 | 2.177 | 0.2187 |

*p*-value adjustment: tukey method for varying family sizes. $R^2$=0.66

Table S9: Estimated Marginal Means ($\log_{10}$) of 16S rRNA Genome copy numbers over time during *in vitro* fermentation of **inulin** with pig feces inoculum. Pairwise comparisons between incubation times.

| contrast | estimate | se | df | t.ratio | p.value |
|---|---|---|---|---|---|
| incub_time0 - incub_time2 | nonEst | NA | NA | NA | NA |
| incub_time0 - incub_time4 | nonEst | NA | NA | NA | NA |
| incub_time0 - incub_time8 | nonEst | NA | NA | NA | NA |
| incub_time0 - incub_time24 | nonEst | NA | NA | NA | NA |
| incub_time2 - incub_time4 | -0.4537 | 0.404 | 27 | -1.122 | 0.6794 |
| incub_time2 - incub_time8 | -1.0432 | 0.404 | 27 | -2.580 | 0.0700 |
| incub_time2 - incub_time24 | -0.2483 | 0.404 | 27 | -0.614 | 0.9267 |
| incub_time4 - incub_time8 | -0.5895 | 0.404 | 27 | -1.458 | 0.4760 |
| incub_time4 - incub_time24 | 0.2054 | 0.404 | 27 | 0.508 | 0.9565 |
| incub_time8 - incub_time24 | 0.7949 | 0.404 | 27 | 1.966 | 0.2258 |

*p*-value adjustment: tukey method for varying family sizes. $R^2$=0.66

**Table S10:** Estimated Marginal Means ($\log_{10}$) of 16S rRNA Genome copy numbers over time during *in vitro* fermentation of **potato starch** with pig feces inoculum. Pairwise comparisons between incubation times.

| contrast | estimate | se | df | t.ratio | p.value |
|---|---|---|---|---|---|
| incub_time0 - incub_time2 | nonEst | NA | NA | NA | NA |
| incub_time0 - incub_time4 | nonEst | NA | NA | NA | NA |
| incub_time0 - incub_time8 | nonEst | NA | NA | NA | NA |
| incub_time0 - incub_time24 | nonEst | NA | NA | NA | NA |
| incub_time2 - incub_time4 | 0.4087 | 0.404 | 27 | 1.011 | 0.7446 |
| incub_time2 - incub_time8 | -0.5520 | 0.404 | 27 | -1.365 | 0.5312 |
| incub_time2 - incub_time24 | -0.3322 | 0.404 | 27 | -0.822 | 0.8437 |
| incub_time4 - incub_time8 | -0.9607 | 0.404 | 27 | -2.376 | 0.1061 |
| incub_time4 - incub_time24 | -0.7410 | 0.404 | 27 | -1.832 | 0.2807 |
| incub_time8 - incub_time24 | 0.2198 | 0.404 | 27 | 0.543 | 0.9475 |

*p*-value adjustment: tukey method for varying family sizes. $R^2$=0.66

**Table S11:** Estimated Marginal Means ($\log_{10}$) of 16S rRNA Genome copy numbers over time during *in vitro* fermentation of **sugar beet pulp** with pig feces inoculum. Pairwise comparisons between incubation times.

| contrast | estimate | se | df | t.ratio | p.value |
|---|---|---|---|---|---|
| incub_time0 - incub_time2 | nonEst | NA | NA | NA | NA |
| incub_time0 - incub_time4 | nonEst | NA | NA | NA | NA |
| incub_time0 - incub_time8 | nonEst | NA | NA | NA | NA |
| incub_time0 - incub_time24 | nonEst | NA | NA | NA | NA |
| incub_time2 - incub_time4 | -0.3371 | 0.404 | 27 | -0.834 | 0.8380 |
| incub_time2 - incub_time8 | -0.4792 | 0.404 | 27 | -1.185 | 0.6413 |
| incub_time2 - incub_time24 | -0.1405 | 0.404 | 27 | -0.348 | 0.9853 |
| incub_time4 - incub_time8 | -0.1421 | 0.404 | 27 | -0.351 | 0.9848 |
| incub_time4 - incub_time24 | 0.1966 | 0.404 | 27 | 0.486 | 0.9615 |
| incub_time8 - incub_time24 | 0.3386 | 0.404 | 27 | 0.837 | 0.8362 |

*p*-value adjustment: tukey method for varying family sizes. $R^2$=0.66

**Table S12:** Estimated Marginal Means ($\log_{10}$) of 16S rRNA Genome copy numbers over time during *in vitro* fermentation of **wheat straw** with pig feces inoculum. Pairwise comparisons between incubation times.

| contrast | estimate | se | df | t.ratio | p.value |
|---|---|---|---|---|---|
| incub_time0 - incub_time2 | nonEst | NA | NA | NA | NA |
| incub_time0 - incub_time4 | nonEst | NA | NA | NA | NA |
| incub_time0 - incub_time8 | nonEst | NA | NA | NA | NA |
| incub_time0 - incub_time24 | nonEst | NA | NA | NA | NA |
| incub_time2 - incub_time4 | -0.0279 | 0.404 | 27 | -0.069 | 0.9999 |
| incub_time2 - incub_time8 | -0.5808 | 0.404 | 27 | -1.436 | 0.4886 |
| incub_time2 - incub_time24 | 0.1604 | 0.404 | 27 | 0.397 | 0.9784 |
| incub_time4 - incub_time8 | -0.5528 | 0.404 | 27 | -1.367 | 0.5300 |
| incub_time4 - incub_time24 | 0.1883 | 0.404 | 27 | 0.466 | 0.9659 |
| incub_time8 - incub_time24 | 0.7412 | 0.404 | 27 | 1.833 | 0.2805 |

*p*-value adjustment: tukey method for varying family sizes. $R^2$=0.66

**Comparison of different substrates:** To evaluate the effects of different substrates on bacterial genome copy numbers, estimated marginal means (EMMs) for substrate were computed, stratified by incubation time. A linear model was fitted to $\log_{10}$-transformed genome copy numbers, including incubation time, substrate, and their interaction as fixed effects. Pairwise comparisons between substrate levels were performed. No significant differences were found.

Table S13: Pairwise comparisons of substrate effects on $\log_{10}$-transformed genome copy numbers at 0 hours of incubation.

| contrast | estimate | se | df | t.ratio | p.value |
|---|---|---|---|---|---|
| Feces - Inoculum | 0.77616 | 0.330 | 27 | 2.351 | 0.0657 |
| Feces - Inulin | nonEst | NA | NA | NA | NA |
| Feces - (pool_feces+buffer) | 0.53354 | 0.330 | 27 | 1.616 | 0.2563 |
| Feces - Potato starch | nonEst | NA | NA | NA | NA |
| Feces - Sugar beet pulp | nonEst | NA | NA | NA | NA |
| Feces - Wheat straw | nonEst | NA | NA | NA | NA |
| Inoculum - Inulin | nonEst | NA | NA | NA | NA |
| Inoculum - (pool_feces+buffer) | -0.24262 | 0.404 | 27 | -0.600 | 0.8213 |
| Inoculum - Potato starch | nonEst | NA | NA | NA | NA |
| Inoculum - Sugar beet pulp | nonEst | NA | NA | NA | NA |
| Inoculum - Wheat straw | nonEst | NA | NA | NA | NA |
| Inulin - (pool_feces+buffer) | nonEst | NA | NA | NA | NA |
| Inulin - Potato starch | nonEst | NA | NA | NA | NA |
| Inulin - Sugar beet pulp | nonEst | NA | NA | NA | NA |
| Inulin - Wheat straw | nonEst | NA | NA | NA | NA |
| (pool_feces+buffer) - Potato starch | nonEst | NA | NA | NA | NA |
| (pool_feces+buffer) - Sugar beet pulp | nonEst | NA | NA | NA | NA |
| (pool_feces+buffer) - Wheat straw | nonEst | NA | NA | NA | NA |
| Potato starch - Sugar beet pulp | nonEst | NA | NA | NA | NA |
| Potato starch - Wheat straw | nonEst | NA | NA | NA | NA |

*p*-value adjustment: tukey method for varying family sizes. $R^2$=0.66

**Table S14**: Pairwise comparisons of substrate effects on $log_{10}$-transformed genome copy numbers at **2 hours** of incubation

| contrast | estimate | se | df | t.ratio | p.value |
|---|---|---|---|---|---|
| Feces - Inoculum | nonEst | NA | NA | NA | NA |
| Feces - Inulin | nonEst | NA | NA | NA | NA |
| Feces - (pool_feces+buffer) | nonEst | NA | NA | NA | NA |
| Feces - Potato starch | nonEst | NA | NA | NA | NA |
| Feces - Sugar beet pulp | nonEst | NA | NA | NA | NA |
| Feces - Wheat straw | nonEst | NA | NA | NA | NA |
| Inoculum - Inulin | 0.73325 | 0.404 | 27 | 1.813 | 0.3870 |
| Inoculum - (pool_feces+buffer) | nonEst | NA | NA | NA | NA |
| Inoculum - Potato starch | 0.36355 | 0.404 | 27 | 0.899 | 0.8947 |
| Inoculum - Sugar beet pulp | 0.68806 | 0.404 | 27 | 1.702 | 0.4498 |
| Inoculum - Wheat straw | 0.61894 | 0.404 | 27 | 1.531 | 0.5525 |
| Inulin - (pool_feces+buffer) | nonEst | NA | NA | NA | NA |
| Inulin - Potato starch | -0.36969 | 0.404 | 27 | -0.914 | 0.8889 |
| Inulin - Sugar beet pulp | -0.04519 | 0.404 | 27 | -0.112 | 1.0000 |
| Inulin - Wheat straw | -0.11431 | 0.404 | 27 | -0.283 | 0.9985 |
| (pool_feces+buffer) - Potato starch | nonEst | NA | NA | NA | NA |
| (pool_feces+buffer) - Sugar beet pulp | nonEst | NA | NA | NA | NA |
| (pool_feces+buffer) - Wheat straw | nonEst | NA | NA | NA | NA |
| Potato starch - Sugar beet pulp | 0.32451 | 0.404 | 27 | 0.802 | 0.9276 |
| Potato starch - Wheat straw | 0.25538 | 0.404 | 27 | 0.632 | 0.9686 |
| Sugar beet pulp - Wheat straw | -0.06912 | 0.404 | 27 | -0.171 | 0.9998 |

*p*-value adjustment: tukey method for varying family sizes. $R^2$=0.66

**Table S15**: Pairwise comparisons of substrate effects on $log_{10}$-transformed genome copy numbers at **4 hours** of incubation

| contrast | estimate | se | df | t.ratio | p.value |
|---|---|---|---|---|---|
| Feces - Inoculum | nonEst | NA | NA | NA | NA |
| Feces - Inulin | nonEst | NA | NA | NA | NA |
| Feces - (pool_feces+buffer) | nonEst | NA | NA | NA | NA |
| Feces - Potato starch | nonEst | NA | NA | NA | NA |
| Feces - Sugar beet pulp | nonEst | NA | NA | NA | NA |
| Feces - Wheat straw | nonEst | NA | NA | NA | NA |
| Inoculum - Inulin | -0.06551 | 0.404 | 27 | -0.162 | 0.9998 |
| Inoculum - (pool_feces+buffer) | nonEst | NA | NA | NA | NA |
| Inoculum - Potato starch | 0.42723 | 0.404 | 27 | 1.057 | 0.8266 |
| Inoculum - Sugar beet pulp | 0.00589 | 0.404 | 27 | 0.015 | 1.0000 |
| Inoculum - Wheat straw | 0.24598 | 0.404 | 27 | 0.608 | 0.9726 |
| Inulin - (pool_feces+buffer) | nonEst | NA | NA | NA | NA |
| Inulin - Potato starch | 0.49274 | 0.404 | 27 | 1.219 | 0.7409 |
| Inulin - Sugar beet pulp | 0.07140 | 0.404 | 27 | 0.177 | 0.9998 |
| Inulin - Wheat straw | 0.31148 | 0.404 | 27 | 0.770 | 0.9370 |
| (pool_feces+buffer) - Potato starch | nonEst | NA | NA | NA | NA |
| (pool_feces+buffer) - Sugar beet pulp | nonEst | NA | NA | NA | NA |
| (pool_feces+buffer) - Wheat straw | nonEst | NA | NA | NA | NA |
| Potato starch - Sugar beet pulp | -0.42134 | 0.404 | 27 | -1.042 | 0.8336 |
| Potato starch - Wheat straw | -0.18125 | 0.404 | 27 | -0.448 | 0.9912 |
| Sugar beet pulp - Wheat straw | 0.24009 | 0.404 | 27 | 0.594 | 0.9749 |

*p*-value adjustment: tukey method for varying family sizes. $R^2$=0.66

**Table S16**: Pairwise comparisons of substrate effects on $log_{10}$-transformed genome copy numbers at **8 hours** of incubation

| contrast | estimate | se | df | t.ratio | p.value |
|---|---|---|---|---|---|
| Feces - Inoculum | nonEst | NA | NA | NA | NA |
| Feces - Inulin | nonEst | NA | NA | NA | NA |
| Feces - (pool_feces+buffer) | nonEst | NA | NA | NA | NA |
| Feces - Potato starch | nonEst | NA | NA | NA | NA |
| Feces - Sugar beet pulp | nonEst | NA | NA | NA | NA |
| Feces - Wheat straw | nonEst | NA | NA | NA | NA |
| Inoculum - Inulin | -0.83854 | 0.404 | 27 | -2.074 | 0.2603 |
| Inoculum - (pool_feces+buffer) | nonEst | NA | NA | NA | NA |
| Inoculum - Potato starch | -0.71707 | 0.404 | 27 | -1.773 | 0.4090 |
| Inoculum - Sugar beet pulp | -0.31974 | 0.404 | 27 | -0.791 | 0.9311 |
| Inoculum - Wheat straw | -0.49044 | 0.404 | 27 | -1.213 | 0.7441 |
| Inulin - (pool_feces+buffer) | nonEst | NA | NA | NA | NA |
| Inulin - Potato starch | 0.12148 | 0.404 | 27 | 0.300 | 0.9981 |
| Inulin - Sugar beet pulp | 0.51881 | 0.404 | 27 | 1.283 | 0.7035 |
| Inulin - Wheat straw | 0.34810 | 0.404 | 27 | 0.861 | 0.9086 |
| (pool_feces+buffer) - Potato starch | nonEst | NA | NA | NA | NA |
| (pool_feces+buffer) - Sugar beet pulp | nonEst | NA | NA | NA | NA |
| (pool_feces+buffer) - Wheat straw | nonEst | NA | NA | NA | NA |
| Potato starch - Sugar beet pulp | 0.39733 | 0.404 | 27 | 0.983 | 0.8607 |
| Potato starch - Wheat straw | 0.22663 | 0.404 | 27 | 0.560 | 0.9796 |
| Sugar beet pulp - Wheat straw | -0.17070 | 0.404 | 27 | -0.422 | 0.9930 |

*p*-value adjustment: tukey method for varying family sizes. $R^2$=0.66

**Table S17**: Pairwise comparisons of substrate effects on $log_{10}$-transformed genome copy numbers at **24 hours** of incubation

| contrast | estimate | se | df | t.ratio | p.value |
|---|---|---|---|---|---|
| Feces - Inoculum | nonEst | NA | NA | NA | NA |
| Feces - Inulin | nonEst | NA | NA | NA | NA |
| Feces - (pool_feces+buffer) | nonEst | NA | NA | NA | NA |
| Feces - Potato starch | nonEst | NA | NA | NA | NA |
| Feces - Sugar beet pulp | nonEst | NA | NA | NA | NA |
| Feces - Wheat straw | nonEst | NA | NA | NA | NA |
| Inoculum - Inulin | -0.92404 | 0.404 | 27 | -2.285 | 0.1807 |
| Inoculum - (pool_feces+buffer) | nonEst | NA | NA | NA | NA |
| Inoculum - Potato starch | -1.37767 | 0.404 | 27 | -3.407 | 0.0163 |
| Inoculum - Sugar beet pulp | -0.86147 | 0.404 | 27 | -2.130 | 0.2368 |
| Inoculum - Wheat straw | -0.62963 | 0.404 | 27 | -1.557 | 0.5362 |
| Inulin - (pool_feces+buffer) | nonEst | NA | NA | NA | NA |
| Inulin - Potato starch | -0.45363 | 0.404 | 27 | -1.122 | 0.7937 |
| Inulin - Sugar beet pulp | 0.06258 | 0.404 | 27 | 0.155 | 0.9999 |
| Inulin - Wheat straw | 0.29441 | 0.404 | 27 | 0.728 | 0.9481 |
| (pool_feces+buffer) - Potato starch | nonEst | NA | NA | NA | NA |
| (pool_feces+buffer) - Sugar beet pulp | nonEst | NA | NA | NA | NA |
| (pool_feces+buffer) - Wheat straw | nonEst | NA | NA | NA | NA |
| Potato starch - Sugar beet pulp | 0.51620 | 0.404 | 27 | 1.277 | 0.7073 |
| Potato starch - Wheat straw | 0.74804 | 0.404 | 27 | 1.850 | 0.3674 |
| Sugar beet pulp - Wheat straw | 0.23184 | 0.404 | 27 | 0.573 | 0.9779 |

*p*-value adjustment: tukey method for varying family sizes. $R^2$=0.66
