## Supplementary File 2 for "From Substrate to Short Chain Fatty Acids: An *In Vitro* Investigation of Bacterial Dynamics in the Porcine Colon using the Modified Hohenheim Gas Test"

**Table S3:** Results of pairwise comparisons of **total SCFA** production between substrates at 2, 4, 8, and 24 hours of *in vitro* fermentation, based on two-way ANOVA with substrate and incubation time as fixed factors. Substrate contrasts are grouped by incubation time (2 h, 4 h, 8 h, 24 h)

| Linear Hypotheses | estimate | se | df | t.ratio | p.value |
|---|---|---|---|---|---|
| inoculum - inulin, 2 == 0 | -58.379 | 5.102 | -11.442 | <0.001 | *** |
| inoculum - starch, 2 == 0 | -6.143 | 5.102 | -1.204 | 0.9926 | |
| inoculum - straw, 2 == 0 | -10.704 | 5.102 | -2.098 | 0.6344 | |
| inoculum - sugar beet pulp, 2 == 0 | -40.559 | 5.102 | -7.949 | <0.001 | *** |
| inulin - starch, 2 == 0 | 52.235 | 5.102 | 10.238 | <0.001 | *** |
| inulin - straw, 2 == 0 | 47.675 | 5.102 | 9.344 | <0.001 | *** |
| inulin - sugar beet pulp, 2 == 0 | 17.819 | 5.102 | 3.492 | 0.0453 | * |
| starch - straw, 2 == 0 | -4.56 | 5.102 | -0.894 | 0.9997 | |
| starch - sugar beet pulp, 2 == 0 | -34.416 | 5.102 | -6.745 | <0.001 | *** |
| straw - sugar beet pulp, 2 == 0 | -29.855 | 5.102 | -5.851 | <0.001 | *** |
| inoculum - inulin, 4 == 0 | -93.628 | 5.102 | -18.35 | <0.001 | *** |
| inoculum - starch, 4 == 0 | -11.624 | 5.102 | -2.278 | 0.507 | |
| inoculum - straw, 4 == 0 | -11.819 | 5.102 | -2.316 | 0.4807 | |
| inoculum - sugar beet pulp, 4 == 0 | -40.234 | 5.102 | -7.885 | <0.001 | *** |
| inulin - starch, 4 == 0 | 82.003 | 5.102 | 16.072 | <0.001 | *** |
| inulin - straw, 4 == 0 | 81.808 | 5.102 | 16.033 | <0.001 | *** |
| inulin - sugar beet pulp, 4 == 0 | 53.394 | 5.102 | 10.465 | <0.001 | *** |
| starch - straw, 4 == 0 | -0.195 | 5.102 | -0.038 | 1 | |
| starch - sugar beet pulp, 4 == 0 | -28.609 | 5.102 | -5.607 | <0.001 | *** |
| straw - sugar beet pulp, 4 == 0 | -28.414 | 5.102 | -5.569 | <0.001 | *** |
| inoculum - inulin, 8 == 0 | -98.497 | 5.102 | -19.304 | <0.001 | *** |
| inoculum - starch, 8 == 0 | -68.322 | 5.102 | -13.39 | <0.001 | *** |
| inoculum - straw, 8 == 0 | -29.307 | 5.102 | -5.744 | <0.001 | *** |
| inoculum - sugar beet pulp, 8 == 0 | -59.871 | 5.102 | -11.734 | <0.001 | *** |
| inulin - starch, 8 == 0 | 30.175 | 5.102 | 5.914 | <0.001 | *** |
| inulin - straw, 8 == 0 | 69.19 | 5.102 | 13.561 | <0.001 | *** |
| inulin - sugar beet pulp, 8 == 0 | 38.626 | 5.102 | 7.57 | <0.001 | *** |
| starch - straw, 8 == 0 | 39.015 | 5.102 | 7.647 | <0.001 | *** |
| starch - sugar beet pulp, 8 == 0 | 8.451 | 5.102 | 1.656 | 0.8961 | |
| straw - sugar beet pulp, 8 == 0 | -30.564 | 5.102 | -5.99 | <0.001 | *** |
| inoculum - inulin, 24 == 0 | -100.591 | 3.608 | -27.881 | <0.001 | *** |
| inoculum - starch, 24 == 0 | -87.486 | 3.608 | -24.248 | <0.001 | *** |
| inoculum - straw, 24 == 0 | -23.78 | 3.608 | -6.591 | <0.001 | *** |
| inoculum - sugar beet pulp, 24 == 0 | -93.74 | 3.608 | -25.982 | <0.001 | *** |
| inulin - starch, 24 == 0 | 13.105 | 3.608 | 3.632 | 0.0323 | * |
| inulin - straw, 24 == 0 | 76.811 | 3.608 | 21.29 | <0.001 | *** |
| inulin - sugar beet pulp, 24 == 0 | 6.851 | 3.608 | 1.899 | 0.7683 | |
| starch - straw, 24 == 0 | 63.706 | 3.608 | 17.657 | <0.001 | *** |
| starch - sugar beet pulp, 24 == 0 | -6.255 | 3.608 | -1.734 | 0.8611 | |
| straw - sugar beet pulp, 24 == 0 | -69.96 | 3.608 | -19.391 | <0.001 | *** |

Two-way ANOVA was performed with substrate and incubation time as factors. Adjusted p-values (single-step method) are reported. Adjusted $R^2 = 0.99$. Comparison of substrate levels included. se = standard error of the estimate, df = degrees of freedom. Significance codes: *** $p < 0.001$, ** $p < 0.01$, * $p < 0.05$, · $p < 0.1$

Table S4: Results of pairwise comparisons of **acetic acid** production between substrates at 2, 4, 8, and 24 hours of *in vitro* fermentation, based on two-way ANOVA with substrate and incubation time as fixed factors. Substrate contrasts are grouped by incubation time (2 h, 4 h, 8 h, 24 h)

| Linear Hypotheses | estimate | se | df | t.ratio | p.value |
|---|---|---|---|---|---|
| inoculum - inulin, 2 == 0 | -15.6732 | 2.6382 | -5.941 | < 0.001 | *** |
| inoculum - starch, 2 == 0 | -2.5831 | 2.6382 | -0.979 | 0.9992 | |
| inoculum - straw, 2 == 0 | -5.964 | 2.6382 | -2.261 | 0.51921 | |
| inoculum - sugar beet pulp, 2 == 0 | -21.0181 | 2.6382 | -7.967 | < 0.001 | *** |
| inulin - starch, 2 == 0 | 13.0902 | 2.6382 | 4.962 | < 0.001 | *** |
| inulin - straw, 2 == 0 | 9.7092 | 2.6382 | 3.68 | 0.02848 | * |
| inulin - sugar beet pulp, 2 == 0 | -5.3448 | 2.6382 | -2.026 | 0.68427 | |
| starch - straw, 2 == 0 | -3.381 | 2.6382 | -1.282 | 0.98657 | |
| starch - sugar beet pulp, 2 == 0 | -18.435 | 2.6382 | -6.988 | < 0.001 | *** |
| straw - sugar beet pulp, 2 == 0 | -15.054 | 2.6382 | -5.706 | < 0.001 | *** |
| inoculum - inulin, 4 == 0 | -35.0254 | 2.6382 | -13.276 | < 0.001 | *** |
| inoculum - starch, 4 == 0 | -4.8835 | 2.6382 | -1.851 | 0.79721 | |
| inoculum - straw, 4 == 0 | -6.9069 | 2.6382 | -2.618 | 0.29661 | |
| inoculum - sugar beet pulp, 4 == 0 | -21.8932 | 2.6382 | -8.299 | < 0.001 | *** |
| inulin - starch, 4 == 0 | 30.142 | 2.6382 | 11.425 | < 0.001 | *** |
| inulin - straw, 4 == 0 | 28.1186 | 2.6382 | 10.658 | < 0.001 | *** |
| inulin - sugar beet pulp, 4 == 0 | 13.1322 | 2.6382 | 4.978 | < 0.001 | *** |
| starch - straw, 4 == 0 | -2.0234 | 2.6382 | -0.767 | 0.99996 | |
| starch - sugar beet pulp, 4 == 0 | -17.0097 | 2.6382 | -6.447 | < 0.001 | *** |
| straw - sugar beet pulp, 4 == 0 | -14.9863 | 2.6382 | -5.681 | < 0.001 | *** |
| inoculum - inulin, 8 == 0 | -36.3541 | 2.6382 | -13.78 | < 0.001 | *** |
| inoculum - starch, 8 == 0 | -28.6225 | 2.6382 | -10.849 | < 0.001 | *** |
| inoculum - straw, 8 == 0 | -16.4923 | 2.6382 | -6.251 | < 0.001 | *** |
| inoculum - sugar beet pulp, 8 == 0 | -33.4231 | 2.6382 | -12.669 | < 0.001 | *** |
| inulin - starch, 8 == 0 | 7.7316 | 2.6382 | 2.931 | 0.16202 | |
| inulin - straw, 8 == 0 | 19.8618 | 2.6382 | 7.529 | < 0.001 | *** |
| inulin - sugar beet pulp, 8 == 0 | 2.931 | 2.6382 | 1.111 | 0.99673 | |
| starch - straw, 8 == 0 | 12.1302 | 2.6382 | 4.598 | 0.00242 | ** |
| starch - sugar beet pulp, 8 == 0 | -4.8006 | 2.6382 | -1.82 | 0.81554 | |
| straw - sugar beet pulp, 8 == 0 | -16.9308 | 2.6382 | -6.418 | < 0.001 | *** |
| inoculum - inulin, 24 == 0 | -37.1294 | 1.8655 | -19.903 | < 0.001 | *** |
| inoculum - starch, 24 == 0 | -36.3922 | 1.8655 | -19.508 | < 0.001 | *** |
| inoculum - straw, 24 == 0 | -17.2518 | 1.8655 | -9.248 | < 0.001 | *** |
| inoculum - sugar beet pulp, 24 == 0 | -50.8176 | 1.8655 | -27.241 | < 0.001 | *** |
| inulin - starch, 24 == 0 | 0.7373 | 1.8655 | 0.395 | 1 | |
| inulin - straw, 24 == 0 | 19.8777 | 1.8655 | 10.655 | < 0.001 | *** |
| inulin - sugar beet pulp, 24 == 0 | -13.6882 | 1.8655 | -7.338 | < 0.001 | *** |
| starch - straw, 24 == 0 | 19.1404 | 1.8655 | 10.26 | < 0.001 | *** |
| starch - sugar beet pulp, 24 == 0 | -14.4254 | 1.8655 | -7.733 | < 0.001 | *** |
| straw - sugar beet pulp, 24 == 0 | -33.5659 | 1.8655 | -17.993 | < 0.001 | *** |

Two-way ANOVA was performed with substrate and incubation time as factors. Adjusted p-values (single-step method) are reported. Adjusted $R^2 = 0.99$. Comparison of substrate levels included. se = standard error of the estimate, df = degrees of freedom. Significance codes: *** $p < 0.001$, ** $p < 0.01$, * $p < 0.05$, · $p < 0.1$

Table S5: Results of pairwise comparisons of **propionic acid** production between substrates at 2, 4, 8, and 24 hours of *in vitro* fermentation, based on two-way ANOVA with substrate and incubation time as fixed factors. Substrate contrasts are grouped by incubation time (2 h, 4 h, 8 h, 24 h)

| Linear Hypotheses | estimate | se | df | t.ratio | p.value |
|---|---|---|---|---|---|
| inoculum - inulin, 2 == 0 | -15.671 | 1.3291 | -11.79 | < 0.001 | *** |
| inoculum - starch, 2 == 0 | -1.6675 | 1.3291 | -1.255 | 0.98901 | |
| inoculum - straw, 2 == 0 | -2.6551 | 1.3291 | -1.998 | 0.70381 | |
| inoculum - sugar beet pulp, 2 == 0 | -12.0196 | 1.3291 | -9.043 | < 0.001 | *** |
| inulin - starch, 2 == 0 | 14.0035 | 1.3291 | 10.536 | < 0.001 | *** |
| inulin - straw, 2 == 0 | 13.0159 | 1.3291 | 9.793 | < 0.001 | *** |
| inulin - sugar beet pulp, 2 == 0 | 3.6515 | 1.3291 | 2.747 | 0.23349 | |
| starch - straw, 2 == 0 | -0.9876 | 1.3291 | -0.743 | 0.99997 | |
| starch - sugar beet pulp, 2 == 0 | -10.352 | 1.3291 | -7.789 | < 0.001 | *** |
| straw - sugar beet pulp, 2 == 0 | -9.3644 | 1.3291 | -7.046 | < 0.001 | *** |
| inoculum - inulin, 4 == 0 | -40.2268 | 1.3291 | -30.266 | < 0.001 | *** |
| inoculum - starch, 4 == 0 | -3.5701 | 1.3291 | -2.686 | 0.26204 | |
| inoculum - straw, 4 == 0 | -2.8697 | 1.3291 | -2.159 | 0.59146 | |
| inoculum - sugar beet pulp, 4 == 0 | -12.1562 | 1.3291 | -9.146 | < 0.001 | *** |
| inulin - starch, 4 == 0 | 36.6568 | 1.3291 | 27.58 | < 0.001 | *** |
| inulin - straw, 4 == 0 | 37.3572 | 1.3291 | 28.106 | < 0.001 | *** |
| inulin - sugar beet pulp, 4 == 0 | 28.0706 | 1.3291 | 21.12 | < 0.001 | *** |
| starch - straw, 4 == 0 | 0.7004 | 1.3291 | 0.527 | 1 | |
| starch - sugar beet pulp, 4 == 0 | -8.5862 | 1.3291 | -6.46 | < 0.001 | *** |
| straw - sugar beet pulp, 4 == 0 | -9.2866 | 1.3291 | -6.987 | < 0.001 | *** |
| inoculum - inulin, 8 == 0 | -42.5464 | 1.3291 | -32.011 | < 0.001 | *** |
| inoculum - starch, 8 == 0 | -23.0849 | 1.3291 | -17.368 | < 0.001 | *** |
| inoculum - straw, 8 == 0 | -7.3775 | 1.3291 | -5.551 | < 0.001 | *** |
| inoculum - sugar beet pulp, 8 == 0 | -17.6312 | 1.3291 | -13.265 | < 0.001 | *** |
| inulin - starch, 8 == 0 | 19.4616 | 1.3291 | 14.642 | < 0.001 | *** |
| inulin - straw, 8 == 0 | 35.1689 | 1.3291 | 26.46 | < 0.001 | *** |
| inulin - sugar beet pulp, 8 == 0 | 24.9152 | 1.3291 | 18.746 | < 0.001 | *** |
| starch - straw, 8 == 0 | 15.7074 | 1.3291 | 11.818 | < 0.001 | *** |
| starch - sugar beet pulp, 8 == 0 | 5.4537 | 1.3291 | 4.103 | 0.00949 | ** |
| straw - sugar beet pulp, 8 == 0 | -10.2537 | 1.3291 | -7.715 | < 0.001 | *** |
| inoculum - inulin, 24 == 0 | -42.15 | 0.9398 | -44.848 | < 0.001 | *** |
| inoculum - starch, 24 == 0 | -26.7125 | 0.9398 | -28.423 | < 0.001 | *** |
| inoculum - straw, 24 == 0 | -6.8226 | 0.9398 | -7.259 | < 0.001 | *** |
| inoculum - sugar beet pulp, 24 == 0 | -26.9313 | 0.9398 | -28.655 | < 0.001 | *** |
| inulin - starch, 24 == 0 | 15.4375 | 0.9398 | 16.426 | < 0.001 | *** |
| inulin - straw, 24 == 0 | 35.3274 | 0.9398 | 37.589 | < 0.001 | *** |
| inulin - sugar beet pulp, 24 == 0 | 15.2188 | 0.9398 | 16.193 | < 0.001 | *** |
| starch - straw, 24 == 0 | 19.8899 | 0.9398 | 21.163 | < 0.001 | *** |
| starch - sugar beet pulp, 24 == 0 | -0.2187 | 0.9398 | -0.233 | 1 | |
| straw - sugar beet pulp, 24 == 0 | -20.1087 | 0.9398 | -21.396 | < 0.001 | *** |

Two-way ANOVA was performed with substrate and incubation time as factors. Adjusted p-values (single-step method) are reported. Adjusted $R^2 = 0.99$. Comparison of substrate levels included. se = standard error of the estimate, df = degrees of freedom. Significance codes: *** $p < 0.001$, ** $p < 0.01$, * $p < 0.05$, · $p < 0.1$

Table S6: Results of pairwise comparisons of **butyric acid** production between substrates at 2, 4, 8, and 24 hours of *in vitro* fermentation, based on two-way ANOVA with substrate and incubation time as fixed factors. Substrate contrasts are grouped by incubation time (2 h, 4 h, 8 h, 24 h)

| Linear Hypotheses | estimate | se | df | t.ratio | p.value |
|---|---|---|---|---|---|
| inoculum - inulin, 2 == 0 | -4.6598 | 0.8944 | -5.21 | < 0.001 | *** |
| inoculum - starch, 2 == 0 | -0.6414 | 0.8944 | -0.717 | 0.99998 | |
| inoculum - straw, 2 == 0 | -1.459 | 0.8944 | -1.631 | 0.90617 | |
| inoculum - sugar beet pulp, 2 == 0 | -4.9781 | 0.8944 | -5.566 | < 0.001 | *** |
| inulin - starch, 2 == 0 | 4.0184 | 0.8944 | 4.493 | 0.00333 | ** |
| inulin - straw, 2 == 0 | 3.2008 | 0.8944 | 3.579 | 0.03674 | * |
| inulin - sugar beet pulp, 2 == 0 | -0.3183 | 0.8944 | -0.356 | 1 | |
| starch - straw, 2 == 0 | -0.8176 | 0.8944 | -0.914 | 0.99964 | |
| starch - sugar beet pulp, 2 == 0 | -4.3367 | 0.8944 | -4.849 | 0.00127 | ** |
| straw - sugar beet pulp, 2 == 0 | -3.5191 | 0.8944 | -3.935 | 0.01488 | * |
| inoculum - inulin, 4 == 0 | -12.3065 | 0.8944 | -13.76 | < 0.001 | *** |
| inoculum - starch, 4 == 0 | -2.1538 | 0.8944 | -2.408 | 0.41974 | |
| inoculum - straw, 4 == 0 | -1.363 | 0.8944 | -1.524 | 0.94237 | |
| inoculum - sugar beet pulp, 4 == 0 | -4.2604 | 0.8944 | -4.764 | 0.00162 | ** |
| inulin - starch, 4 == 0 | 10.1526 | 0.8944 | 11.352 | < 0.001 | *** |
| inulin - straw, 4 == 0 | 10.9435 | 0.8944 | 12.236 | < 0.001 | *** |
| inulin - sugar beet pulp, 4 == 0 | 8.0461 | 0.8944 | 8.997 | < 0.001 | *** |
| starch - straw, 4 == 0 | 0.7908 | 0.8944 | 0.884 | 0.99976 | |
| starch - sugar beet pulp, 4 == 0 | -2.1065 | 0.8944 | -2.355 | 0.45455 | |
| straw - sugar beet pulp, 4 == 0 | -2.8973 | 0.8944 | -3.24 | 0.0825 | . |
| inoculum - inulin, 8 == 0 | -13.0919 | 0.8944 | -14.638 | < 0.001 | *** |
| inoculum - starch, 8 == 0 | -15.6572 | 0.8944 | -17.507 | < 0.001 | *** |
| inoculum - straw, 8 == 0 | -3.5779 | 0.8944 | -4.001 | 0.0126 | * |
| inoculum - sugar beet pulp, 8 == 0 | -6.6374 | 0.8944 | -7.421 | < 0.001 | *** |
| inulin - starch, 8 == 0 | -2.5653 | 0.8944 | -2.868 | 0.18409 | |
| inulin - straw, 8 == 0 | 9.514 | 0.8944 | 10.638 | < 0.001 | *** |
| inulin - sugar beet pulp, 8 == 0 | 6.4544 | 0.8944 | 7.217 | < 0.001 | *** |
| starch - straw, 8 == 0 | 12.0793 | 0.8944 | 13.506 | < 0.001 | *** |
| starch - sugar beet pulp, 8 == 0 | 9.0198 | 0.8944 | 10.085 | < 0.001 | *** |
| straw - sugar beet pulp, 8 == 0 | -3.0595 | 0.8944 | -3.421 | 0.05395 | . |
| inoculum - inulin, 24 == 0 | -14.5917 | 0.6324 | -23.073 | < 0.001 | *** |
| inoculum - starch, 24 == 0 | -18.8101 | 0.6324 | -29.744 | < 0.001 | *** |
| inoculum - straw, 24 == 0 | -1.1072 | 0.6324 | -1.751 | 0.8525 | |
| inoculum - sugar beet pulp, 24 == 0 | -11.146 | 0.6324 | -17.625 | < 0.001 | *** |
| inulin - starch, 24 == 0 | -4.2183 | 0.6324 | -6.67 | < 0.001 | *** |
| inulin - straw, 24 == 0 | 13.4846 | 0.6324 | 21.323 | < 0.001 | *** |
| inulin - sugar beet pulp, 24 == 0 | 3.4457 | 0.6324 | 5.449 | < 0.001 | *** |
| starch - straw, 24 == 0 | 17.7029 | 0.6324 | 27.993 | < 0.001 | *** |
| starch - sugar beet pulp, 24 == 0 | 7.6641 | 0.6324 | 12.119 | < 0.001 | *** |
| straw - sugar beet pulp, 24 == 0 | -10.0389 | 0.6324 | -15.874 | < 0.001 | *** |

Two-way ANOVA was performed with substrate and incubation time as factors. Adjusted p-values (single-step method) are reported. Adjusted $R^2 = 0.99$. Comparison of substrate levels included. se = standard error of the estimate, df = degrees of freedom. Significance codes: *** $p < 0.001$, ** $p < 0.01$, * $p < 0.05$, · $p < 0.1$
