## Supplementary File 1 for "From Substrate to Short Chain Fatty Acids: An *In Vitro* Investigation of Bacterial Dynamics in the Porcine Colon using the Modified Hohenheim Gas Test"

**Table S1:** Summary of gas formation [ml/200 mg substrate] during *in vitro* fermentation of different substrates using the modHGT with pig feces at 2, 4, 8, and 24 hours of incubation.

| Substrate | 2h | | 4h | | 8h | | 24h | |
|---|---|---|---|---|---|---|---|---|
| Inoculum | 0.0 | ± 0.0 | 1.2 | ± 0.3 | 1.2 | ± 0.3 | 4.1 | ± 0.2 |
| Inulin | 5.3 | ± 0.8 | 47.1 | ± 2.5 | 55.9 | ± 0.6 | 59.6 | ± 1.1 |
| Potato starch | 0.3 | ± 0.3 | 1.6 | ± 0.5 | 44.4 | ± 0.7 | 62.7 | ± 1.1 |
| Wheat straw | 0.4 | ± 0.3 | 2.0 | ± 0.5 | 3.7 | ± 0.3 | 17.9 | ± 0.4 |
| Sugar beet pulp | 6.4 | ± 0.4 | 18.0 | ± 0.5 | 31.9 | ± 1.0 | 58.2 | ± 0.5 |

Mean values ± standard deviation are shown. Sample sizes per time point: 2 h (n = 10), 4 h (n = 8), 8 h (n = 6), 24 h (n = 4).

**Table S2:** Statistical analysis of gas formation during *in vitro* fermentation of different Substrates with pig feces at 2-, 4-, 8-, and 24-hours incubation using simultaneous tests for general linear hypotheses.

| Linear Hypotheses | Estimate | se | t value | Pr(>\|t\|) | |
|---|---|---|---|---|---|
| Inoculum - Inulin, 2 == 0 | -5.3161 | 0.3559 | -14.938 | < 0.001 | *** |
| Inoculum - Starch, 2 == 0 | -0.271 | 0.3559 | -0.762 | 0.99998 | |
| Inoculum - Straw, 2 == 0 | -0.3854 | 0.3559 | -1.083 | 0.99855 | |
| Inoculum - Sugar Beet Pulp, 2 == 0 | -6.3566 | 0.3559 | -17.862 | < 0.001 | *** |
| Inulin - Starch, 2 == 0 | 5.0451 | 0.3559 | 14.176 | < 0.001 | *** |
| Inulin - Straw, 2 == 0 | 4.9308 | 0.3559 | 13.855 | < 0.001 | *** |
| Inulin - Sugar Beet Pulp, 2 == 0 | -1.0405 | 0.3559 | -2.924 | 0.12404 | |
| Starch - Straw, 2 == 0 | -0.1144 | 0.3559 | -0.321 | 1 | |
| Starch - Sugar Beet Pulp, 2 == 0 | -6.0856 | 0.3559 | -17.1 | < 0.001 | *** |
| Straw - Sugar Beet Pulp, 2 == 0 | -5.9713 | 0.3559 | -16.779 | < 0.001 | *** |
| Inoculum - Inulin, 4 == 0 | -46.5758 | 0.3979 | -117.057 | < 0.001 | *** |
| Inoculum - Starch, 4 == 0 | -1.106 | 0.3979 | -2.78 | 0.17785 | |
| Inoculum - Straw, 4 == 0 | -1.5462 | 0.3979 | -3.886 | 0.00626 | ** |
| Inoculum - Sugar Beet Pulp, 4 == 0 | -17.5391 | 0.3979 | -44.08 | < 0.001 | *** |
| Inulin - Starch, 4 == 0 | 45.4698 | 0.3979 | 114.277 | < 0.001 | *** |
| Inulin - Straw, 4 == 0 | 45.0297 | 0.3979 | 113.171 | < 0.001 | *** |
| Inulin - Sugar Beet Pulp, 4 == 0 | 29.0368 | 0.3979 | 72.977 | < 0.001 | *** |
| Starch - Straw, 4 == 0 | -0.4402 | 0.3979 | -1.106 | 0.99815 | |
| Starch - Sugar Beet Pulp, 4 == 0 | -16.4331 | 0.3979 | -41.301 | < 0.001 | *** |
| Straw - Sugar Beet Pulp, 4 == 0 | -15.9929 | 0.3979 | -40.194 | < 0.001 | *** |
| Inoculum - Inulin, 8 == 0 | -54.6443 | 0.4594 | -118.936 | < 0.001 | *** |
| Inoculum - Starch, 8 == 0 | -43.1941 | 0.4594 | -94.014 | < 0.001 | *** |
| Inoculum - Straw, 8 == 0 | -2.4966 | 0.4594 | -5.434 | < 0.001 | *** |
| Inoculum - Sugar Beet Pulp, 8 == 0 | -30.6635 | 0.4594 | -66.74 | < 0.001 | *** |
| Inulin - Starch, 8 == 0 | 11.4501 | 0.4594 | 24.922 | < 0.001 | *** |
| Inulin - Straw, 8 == 0 | 52.1477 | 0.4594 | 113.502 | < 0.001 | *** |
| Inulin - Sugar Beet Pulp, 8 == 0 | 23.9808 | 0.4594 | 52.195 | < 0.001 | *** |
| Starch - Straw, 8 == 0 | 40.6975 | 0.4594 | 88.58 | < 0.001 | *** |
| Starch - Sugar Beet Pulp, 8 == 0 | 12.5306 | 0.4594 | 27.273 | < 0.001 | *** |
| Straw - Sugar Beet Pulp, 8 == 0 | -28.1669 | 0.4594 | -61.307 | < 0.001 | *** |
| Inoculum - Inulin, 24 == 0 | -55.5241 | 0.5627 | -98.674 | < 0.001 | *** |
| Inoculum - Starch, 24 == 0 | -58.5809 | 0.5627 | -104.107 | < 0.001 | *** |
| Inoculum - Straw, 24 == 0 | -13.7611 | 0.5627 | -24.455 | < 0.001 | *** |
| Inoculum - Sugar Beet Pulp, 24 == 0 | -54.0768 | 0.5627 | -96.102 | < 0.001 | *** |
| Inulin - Starch, 24 == 0 | -3.0568 | 0.5627 | -5.432 | < 0.001 | *** |
| Inulin - Straw, 24 == 0 | 41.763 | 0.5627 | 74.219 | < 0.001 | *** |
| Inulin - Sugar Beet Pulp, 24 == 0 | 1.4473 | 0.5627 | 2.572 | 0.28432 | |
| Starch - Straw, 24 == 0 | 44.8198 | 0.5627 | 79.651 | < 0.001 | *** |
| Starch - Sugar Beet Pulp, 24 == 0 | 4.5041 | 0.5627 | 8.004 | < 0.001 | *** |
| Straw - Sugar Beet Pulp, 24 == 0 | -40.3157 | 0.5627 | -71.647 | < 0.001 | *** |

*p*-values (single-step method) are reported. Comparison of substrate levels included. Adjusted$R^2$ = 0.90. se = standard error.
Significance codes: *** p < 0.001, ** p < 0.01, * p < 0.05, · p < 0.1
